# Rescue of non-healing, degenerative salivary glands by cholinergic-calcium signaling

**DOI:** 10.1101/2024.12.31.630834

**Authors:** Jianlong Li, Bo Sun, Li Xuan Tan, Nathan Griffin, Seyyed Vahid Niknezhad, Chieh Yu, Lionel Berthoin, Noel Cruz-Pacheco, Seayar Mohabbat, Hanan Sinada, Yael Efraim, Feeling Yu Ting Chen, Luye An, Eliza A. Gaylord, Chelsey S. Bahney, Isabelle M.A. Lombaert, Sarah M. Knox

**Author notes:** These authors contributed equally (Jianlong Li, Bo Sun, Li Xuan Tan). Co–senior authors (Isabelle M.A. Lombaert, Sarah M. Knox).

## Abstract

Chronic degenerative wounds are often deemed irreparable, directing research efforts to focus predominantly on acute tissue injury regeneration while leaving endogenous repair mechanisms for chronically damaged tissues largely unexplored. In this study, we demonstrate that non-healing, severely degenerated salivary gland tissues can be fundamentally restored through first-line treatment with muscarinic agonists. This approach rescues tissue structure and function, returning it to a homeostatic-like state, and reactivates endogenous regeneration processes to drive new cell expansion that persists for months post-treatment. Furthermore, neuromimetic activation profoundly depletes radiation-induced DNA damage and re-establishes the nerve-acinar relationship, ultimately restoring the tissues physiological capacity to maintain homeostasis, even in the absence of treatment. We show that full recovery of organ function, comparable to uninjured controls, is primarily mediated by the re-differentiation of aberrantly de-differentiated epithelial acinar cells and the restoration of mitochondrial function via a muscarinic-calcium signaling pathway. These findings challenge the prevailing notion that chronic organ degeneration is irreversible and propose a readily testable therapeutic strategy for epithelial restoration with potential applications across a spectrum of chronic injuries.

## INTRODUCTION

Chronic injury, characterized by tissue degeneration and destruction, is widely regarded as irreversible ^1–4^. This notion has driven extensive studies into regenerative strategies focused primarily on targeting acute injury states. Current approaches to treating chronic injury—such as device implantation, stem cell therapy, and organ transplantation—often fail to restore the tissue to its pre-injury homeostatic state, further reinforcing the prevailing dogma that chronic tissue damage is irreversible. In this study, we challenge this paradigm by demonstrating, for the first time, that chronically degenerated, epithelial organs can be healed, offering a transformative perspective on regenerative medicine.

Organ degeneration is a major characteristic of chronic injury, manifesting in conditions such as scleroderma (skin) and interstitial lung disease, as well as in response to external exposures that cause severe physical or genomic damage. A prime example of such an external mediator is ionizing radiation (IR), commonly used in cancer treatment. Off-target exposure of healthy tissues—including lung, skin, retina, and glandular organs—to IR often leads to widespread structural and functional tissue decline ^1,5–7^. The acute effects of IR exposure, occurring within days to weeks, can trigger endogenous repair mechanisms that may partially restore organ function ^8^. However, this is followed by chronic tissue degeneration which occurs over months to years, and correlates with partial or complete non-healing tissue dysfunction and destruction ^8–10^. To date, it remains unknown whether chronically injured epithelial organs —such as the lung, intestine, or skin—or tissues affected by other injury or disease states, can be restored once they enter the degenerative phase or remains responsive to regenerative cues after establishing the chronic, non-healing, degenerative state. The submandibular salivary gland (SG) represents an epithelial organ highly susceptible to IR. Inadvertent exposure during the treatment of head and neck cancer routinely leads to progressive tissue destruction and organ failure over a 2-3 month period ^8,11–14^, making the IR SG an ideal platform to study chronic degeneration and potential misconceptions around its limited repair ability.

There is widespread consensus that epithelial stem cell processes mediated by proliferation, differentiation, and interactions with the niche are essential to repair. In fact, clinical neuromimetic treatment suggest that neurogenesis can be activated under various neurodegenerative conditions, such as Alzheimer and epilepsy, by activating neuronal stem cells. It has also been postulated that a non-neuronal neuromimetic system can influence epithelial stem cell fate in various organs (e.g., skin, hair follicle, and intestine), governing a positive involvement in the repair of acute injury models^8,12,15^. However, the appropriate control of epithelial stem cells and their niches by neuromimetics under non-healing, degenerative, conditions have not been explored yet.

Using a series of in vivo and ex vivo murine studies, we reveal that non-healing, chronically degenerated, SGs remain responsive to a neuromimetic treatment. Such first-line treatment effectively restore the tissue to a homeostatic-like state-despite previous belief that such a deteriorated organ is unable to enter a repair program. Remarkably, the restored SGs closely mimic a healthy organ for months after the end of treatment and continue to be responsive to treatments until the homeostatic physiological state is endogenously reestablished, demonstrating both the robustness and sustainability of this regenerative program. These findings challenge the long-standing notion that chronic organ degeneration is irreversible and offer novel therapeutic strategies for other chronically damaged epithelial tissues.

## RESULTS

### Partially destroyed SGs can be functionally regenerated to mimic the uninjured organ

Previous studies by us and others have demonstrated that acutely injured organs (<14 days post-IR) can be regenerated by an array of treatments ^8,16–18^. However, such an outcome has not been reported for chronically injured tissue undergoing degeneration and that no longer of self-healing capacity, likely due to the assumption that chronically damaged epithelial organs, such as intestine, lung, skin and SG, are unresponsive to regenerative cues. To explore the potential for reversal of degeneration as opposed to rescue of acute injury, we focused on the responsiveness of a clinically relevant degenerating exocrine tissue, the irradiated (IR) SG. In this organ, IR results in the chronic loss of acinar cells over time that occurs in conjunction with significant alterations in the tissue microenvironment, including chronic inflammation, aberrant innervation and vascularization (Figure 1A ^8^).

**Figure 1.**
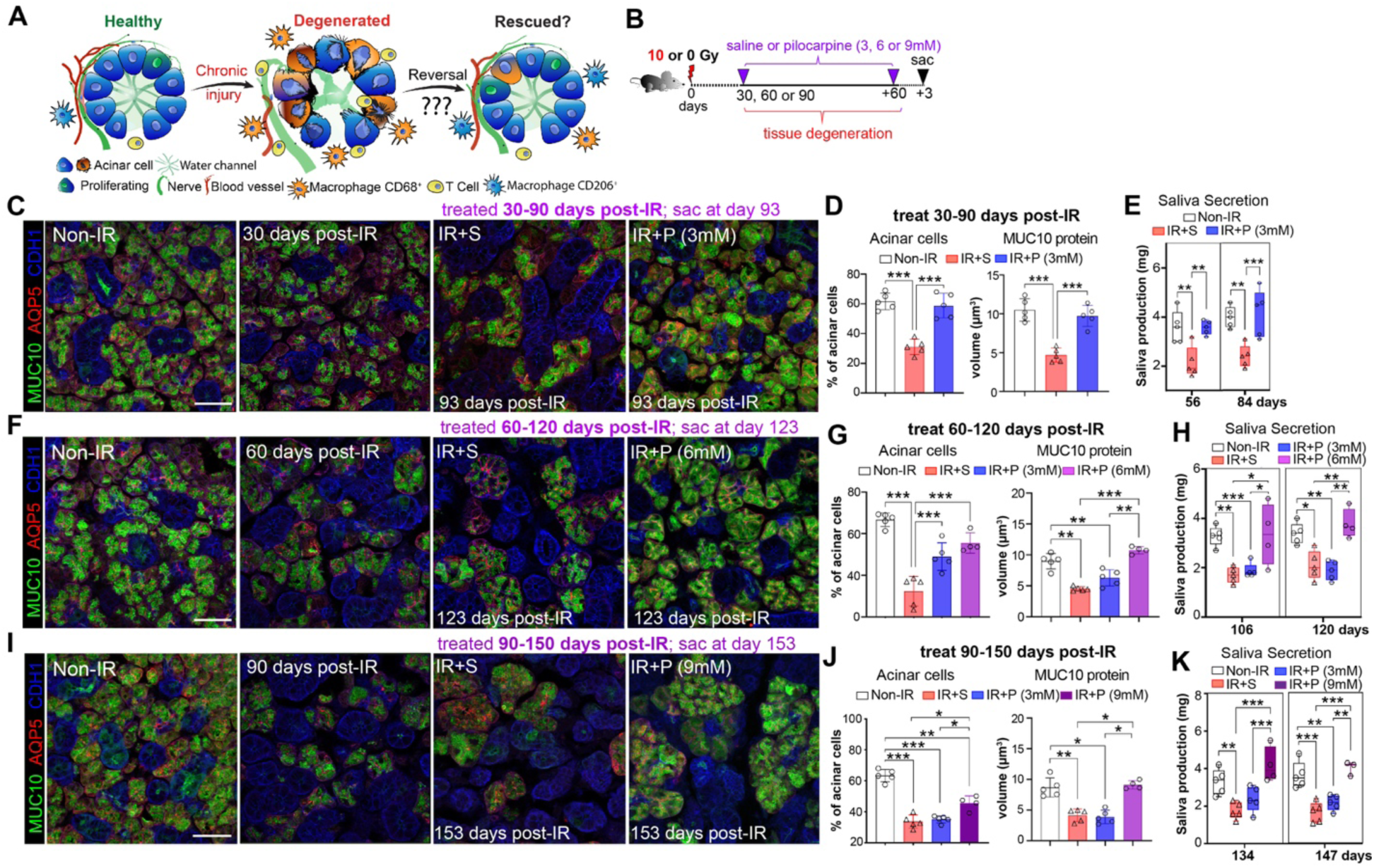
Partially destroyed SGs can be functionally regenerated to mimic the uninjured organ. **A.** Schematic of question being tested, that is, can the destruction of SGs in response to radiation be reversed. **B.** Treatment regimen. **C-E.** Morphological (C-D) and functional (E) analyses of non-IR, IR+S, and IR+P (3 mM) SGs treated from 30-90 days post-IR. Immunostaining (C) and quantification (D) of AQP5+ acinar cells (red) and MUC10 protein (green) at 93-days post-IR. Physiological saliva output at 56- and 84-days post-IR (E). **F-H**. Morphological (F-G) and functional (H) analysis of non-IR, IR+ S, and IR+P (6 mM) SG treated from 60-120 days post-IR. Immunostaining (F) and quantification (G) of AQP5+ acinar cells (red) and MUC10 protein (green) at 123-days post-IR. Physiological saliva output at 106 and 120 days post-IR (H). **I-K.** Morphological (I-J) and functional (K) analysis of non-IR, IR+ S, and IR+P (9 mM) SG treated from 90-150 days post-IR. Immunostaining (I) and quantification (J) of AQP5+ acinar cells (red) and MUC10 protein (green) at 153-days post-IR. Physiological saliva output at 134 and 147 days post-IR (K). Mean±SD. *, *P*<0.05. **, *P*<0.01. ***, *P*<0.001. Scale bar in C, F, and I is 50 µm.

**Figure S1.**
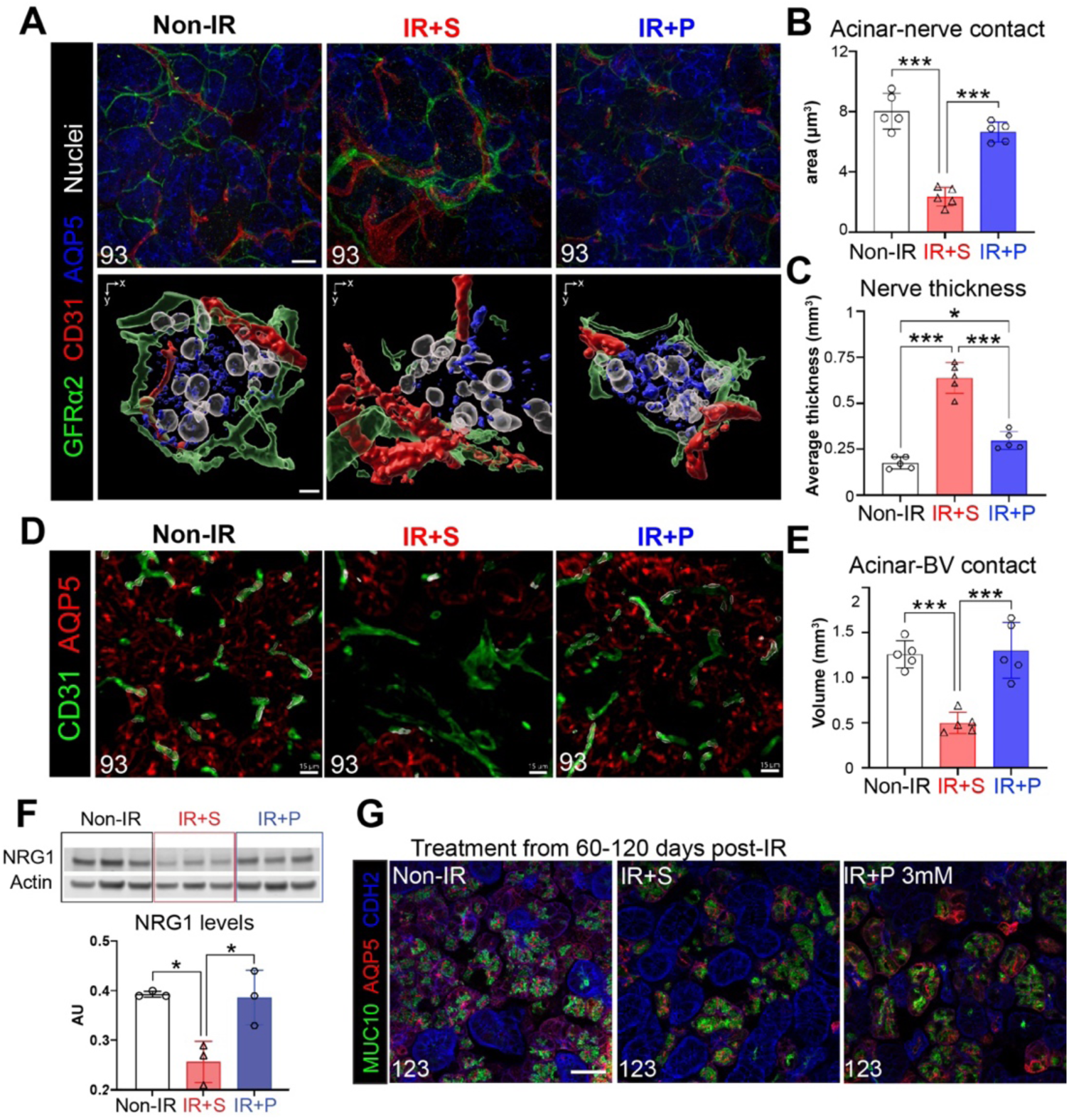
Partially destroyed SGs can be functionally regenerated to mimic the uninjured organ, related to Figure 1. A-C. Immunofluorescent imaging (upper panel) and 3D reconstruction (IMARIS, lower panel) of the acinar (AQP5)-nerve (GFRα2)-capillary (CD31) interactions in SGs of different treatment groups at 93 days post-IR (treatment at 30-90 days post-IR). Quantification of acinar-nerve interactions (B) and large nerve fibers (E). **D-E.** Immunofluorescent imaging (D) and quantification (E) of blood vessels within the SGs of the different experimental groups at 93 days post-IR (treatment at 30-90 days post-IR). **F.** Protein expression and quantification of NRG1 and actin in SG samples at 93 days post-IR. AU, arbitrary units, normalized to actin. **G.** Images of MUC10^+^AQP5^+^ acinar cells of SGs treated 60-120 days post-IR with 3 mM pilocarpine (IR+ P, 3 mM). Tissue was analyzed at 123 days post-IR. Scale bar in A upper panel and G are 20 µm, A lower panel is 5 µm and D is 15 µm. Mean±SD. *, *P*<0.05. ***, *P*<0.001.

To determine whether chronically degenerating IR SGs remain responsive to a regenerative agent, such as a muscarinic mimetic used for acute injury ^8^, we utilized a treatment plan in which mice were given a single 10 Gy dose of IR (equivalent to daily doses to reach 10 Gy^19^) and then injected with the muscarinic agonist pilocarpine (P, 3 mM) or saline (S, IR control), 4 times per week from 30- to 90-days post-IR, with outcomes compared to age matched non-IR controls (Figure 1B). This timeline encompasses the degenerative phase that leads from mild (30 days) to severe (90 days) acinar disruption and destruction ^8,10,20^. Remarkably, we found chronically damaged SGs were highly responsive to pilocarpine treatment (IR+P 3mM), as shown by the extensive recovery of AQP5^+^ acinar cells and levels of the secreted acinar protein MUC10 at 93 days post-IR (3 days after the final injection of pilocarpine) as compared to saline-treated controls (IR+S), effectively mimicking non-IR SGs (Figure 1C-D). Furthermore, pilocarpine treatment completely restored secretory function to that of non-IR tissues (Figure 1E), suggesting functional innervation and vascularization of acini was restored. This possibility was confirmed through immunofluorescent imaging of functional parasympathetic nerves (GFRα2^+^AChE^+^) and capillaries (CD31+) revealing the profound disruption of both circuits in the IR+SG was returned to a homeostatic-like state with pilocarpine (Figure S1A-E). In addition, in support of renewal of acinar-nerve communication, we found levels of NRG1, a protein extensively involved in neuronal-acinar interactions ^21^, to be significantly downregulated in IR+S SGs (46% of non-IR) and returned to those of healthy controls in IR+P SGs (98% of non-IR) (Figure S1F), confirming restoration of the nerve-acinar connections in the rescued tissue.

Next, we questioned whether reversal of degeneration was limited to the early degenerative stage by treating IR mice starting from 60 days post-IR, a time point associated with extensive loss of acinar cells^20^. As expected, SGs from mice treated with saline from 60 to 120 days post-IR showed a profound reduction in AQP5^+^ acinar cells and MUC10 expression by day 123 (Figure 1F-G), with glandular destruction comparable to that observed in IR+S SGs treated from 30 to 90 days (Figure 1C). However, despite pilocarpine promoting acinar cell regeneration (Figure 1F-G, AQP5+ cells), MUC10 protein expression and saliva output remained significantly depleted by 123 days post-IR, matching saline-treated controls (Figure 1F-H). Given this outcome suggested degenerating acinar cells were responsive to muscarinic agonism but not functionally reparative, we tested whether this was due to an insufficient dose of pilocarpine. Strikingly, increasing the concentration of pilocarpine from 3 to 6 mM was sufficient to restore both SG structure and secretory function to non-IR SG levels (Figure 1G-H), demonstrating that a robust reversal of degenerative processes is possible at this time point but requires a higher drug dose.

Building on this finding, we next tested whether SGs at 90 days post-IR — a stage characterized by profound and stable acinar cell destruction — could also be restored by administering an increased dose of 9 mM pilocarpine from 90 to 150 days post-IR. By 153 days post-IR, pilocarpine treatment significantly increased the number of acinar cells (40% above IR+S controls) and restored MUC10 expression to non-IR levels (Figure 1I-J). Remarkably, saliva secretion was also rescued, with levels matching those of non-IR SGs (Figure 1K). These results demonstrate that increasing the treatment dose is sufficient to reestablish normal glandular structure and function, even at advanced stages of chronic degeneration.

Together, these findings reveal that chronically damaged and degenerating SGs remain highly responsive to regenerative cues and can undergo extensive functional repair in a dose-dependent manner.

### Reactivated endogenous repair remains active after treatment termination

Key to the success of restorative therapies is for the regenerated organ to sustain functionally intact after the end of treatment. To determine whether active acinar cell replenishment was sustained after treatment termination, we performed genetic lineage tracing of acinar progenitor cells using *Bmi1^CreERT^*;*Rosa26^RFP^* mice^22^. Similar to the pancreas, *Bmi1* marks a subset of acinar cells^23^ that extensively repopulates the acinar compartment by 30 days post Cre-activation under homeostatic conditions (Figure S2A). However, SGs in which Cre is activated 30 days after IR and traced for 30 days (60 days post-IR) show little to no repopulation of acinar cells in comparison to the non-IR controls (Figure S2B). Using this approach, we investigated acinar cell replenishment after reparative treatment by activating Cre 3 days after the final dose of pilocarpine or saline (i.e., at 93-days post-IR) and collecting tissue 30 days later or 123-days post-IR (Figure 2A). At this time point, we found a significant reduction in RFP+ acinar cells in IR+S SGs compared to the non-IR controls (Figure 2B-C), consistent with the degenerative state of the radiation-damaged tissue. In contrast, the number of RFP+ cells in IR+P SGs was almost indistinguishable from non-IR SG (99% of non-IR RFP+ cells, Figure 2B-C), demonstrating that the formation of new cells from acinar progenitors is retained after treatment termination. The sustained homeostatic-like state of the regenerated tissue was further supported by the restoration of AQP5+ acinar cell numbers (Figure 2D-E), MUC10 protein levels (Figure 2D-F), and functional (AChE+) parasympathetic innervation of acini, all of which mirrored non-IR controls (Figure 2G-H). In stark contrast, IR+S SGs displayed thickened nerve fibers often located within the mesenchyme, distant from the epithelium (Figure 2G, arrows), whereas IR+P SGs exhibited thin nerve fibers directly innervating acini, similar to non-IR SGs (Figure 2G), confirming endogenous repair is also occurring at the surrounding niche level.

**Figure 2.**
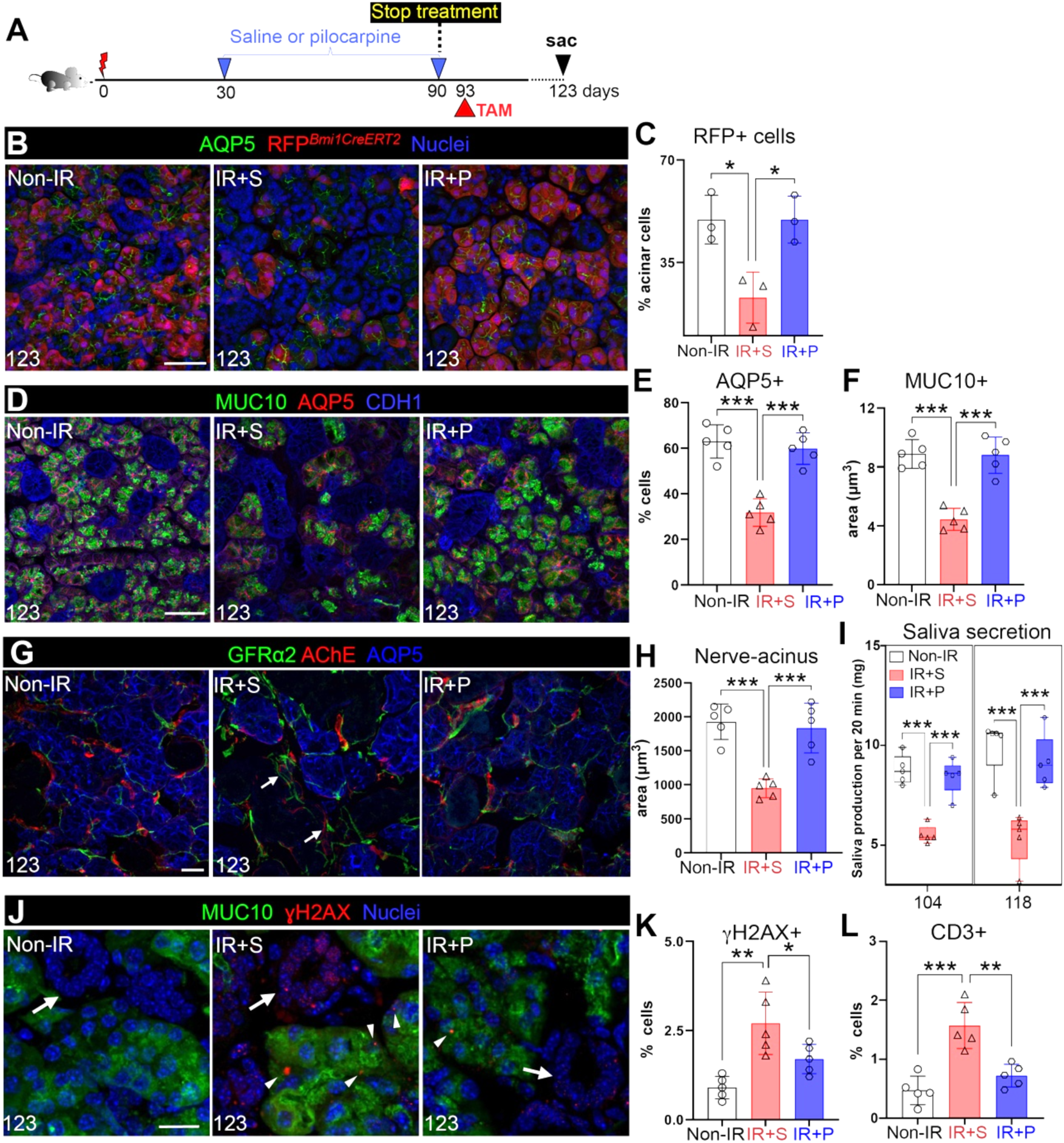
Reactivated endogenous repair remains active after treatment termination. **A.** Schematic of the treatment regimen. **B-C.** Genetic lineage tracing of acinar cells after treatment termination. *Bmi1^CreERT^*^2^*;Rosa26^RFP^* mice were injected with tamoxifen at 93 days post-IR and traced for 30 days. SGs were stained for AQP5 and quantified for the percent of RFP^+^ cells in AQP5^+^ acini. **D-F**. Immunofluorescent analysis of AQP5^+^ and MUC10^+^ acinar cells in non-IR, IR+S and IR+P SGs at 123 days post-IR (D), with subsequent quantification (E-F). **G-H.** Immunofluorescent analysis of GFRα2^+^ and AChE^+^ nerves surrounding acini at day 123 post-IR (E), and quantification of nerve-acinar cell interactions (F). Arrows marks nerves in the mesenchyme. **I**. Physiological saliva secretion measured at day 104 and 118 post-IR. **J-L.** Immunostaining (J) and quantification (K) of MUC10^+^ γH2AX^+^ cells and quantification of CD3^+^ T cells of SGs of the 3 treatment groups. Arrows in J marks ducts, arrow heads mark γH2AX^+^. Mean±SD. *, *P*<0.05. **, *P*<0.01. ***, *P*<0.001. Scale bars in B,D,G are 50 µm and G and J and 20µm.

**Figure S2.**
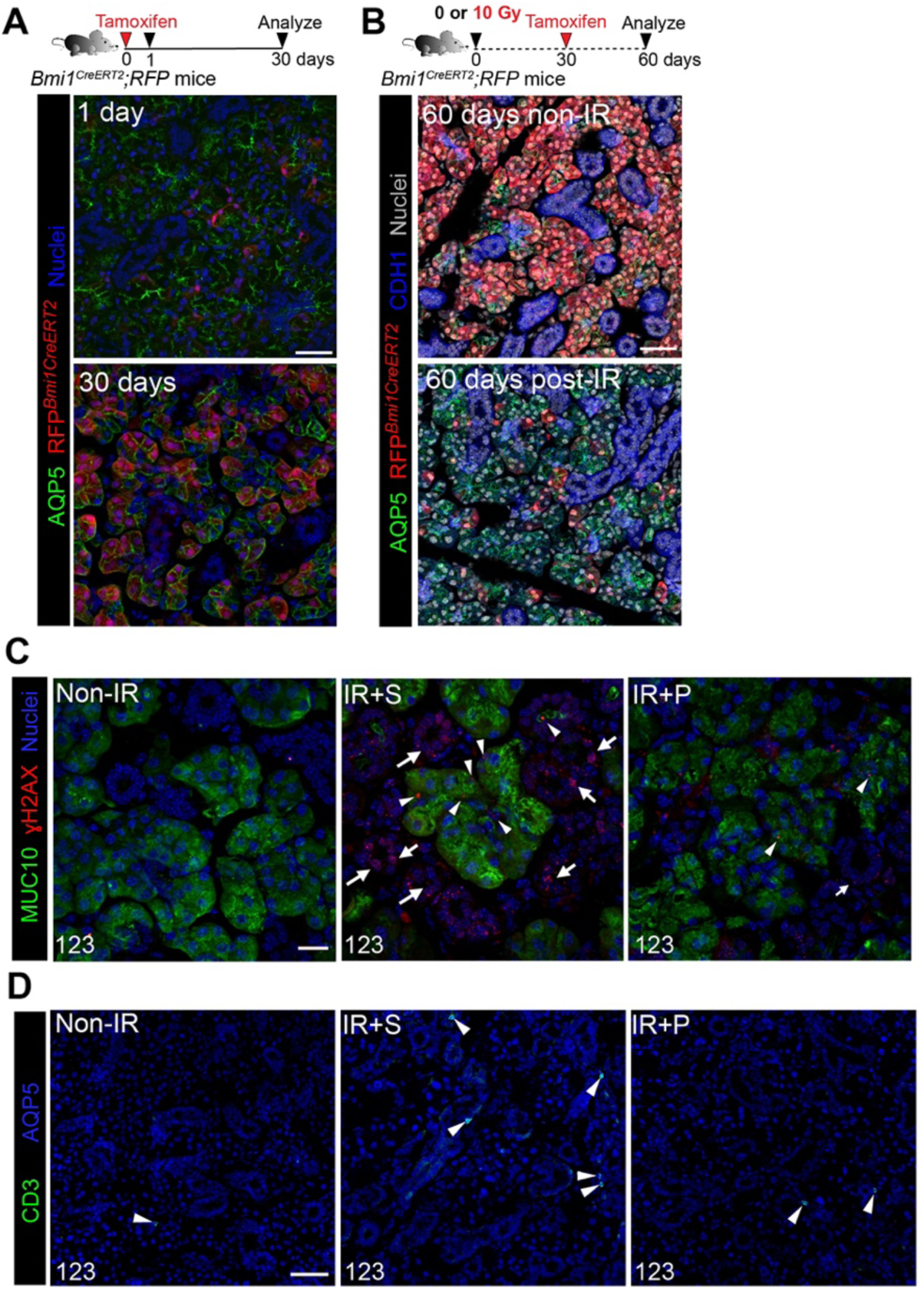
Reactivated endogenous repair remains active after treatment termination, related to Figure 2. **A.** Lineage tracing of *Bmi1^CreERT^*^2^*;Rosa26^RFP^* mice for 1 and 30 days after Cre activation. Mice were injected with tamoxifen at day 0 and immunostained for AQP5 at day 30. **B.** *Bmi1^CreERT^*^2^*;Rosa26^RFP^* mice were treated with 0 or 10 Gy, injected with tamoxifen at 30 days post-IR and immunostained for AQP5 and CDH1 at day 60. **C.** Immunostaining of DNA damage (γH2AX, C) in non-IR, IR+S and IR+P (3 mM, 30-90 days post-IR) at 123-days post-IR (30 days after treatment termination). MUC10 labels acinar cells. Arrowheads mark γH2AX^+^ acinar cells. Arrows mark γH2AX^+^ ducts. **D.** Immunostaining for CD3^+^ T cells in the 3 groups at 123 days post-IR. Arrowheads mark CD3+ cells. Scale bars, 20 μm (A-B), 15 µm (C) and 50 µm (D).

The robustness of tissue maintenance post-treatment was further evidenced by rescued saliva secretion, with IR+P SGs matching the output of non-IR controls (Figure 2I), while IR+S SGs showed a ∼40% reduction (Figure 2I). Additionally, IR+P SGs exhibited significantly lower levels of DNA damage in both acinar cells (γH2AX+ MUC10+ cells, Figure 2J-K and S2C, arrowhead) and non-acinar cells (γH2AX+ MUC10-cells, Figure 2J-K and S2C, arrow), as well as reduced immune cell infiltration compared to IR+S SGs at 123 days post-IR (Figure 2L and S2D, arrowhead). The latter indicates that sustained output of neuromimetic treatment not only rescues organ function via expanding muscarinic-responsive acinar progenitors but has also profound reparative changes in the niche and other non-muscarinic responsive epithelial cell types. This is important since tissue homeostasis requires optimal signaling between nerves and acini (e.g., via the muscarinic pathway) as well as other cell-cell communications. Together, these findings demonstrate that the reversal of chronic damage is not only achievable but also sustained after removal of external neuromimetics, underscoring the potency of this endogenous regenerative approach.

### Functional tissue maintenance is prolonged after the end of treatment, and tissue remains responsive to additional treatments

To determine whether tissue restoration persists or diminishes after treatment cessation, we evaluated the morphological structure and function of SGs over a 5-month period following treatment termination. To clinically mimic a muscarinic agonist with minimal adverse effects, we used cevimeline (CV, 12 mM^8^), which was previously in acute injury studies^8^. Similar to the pilocarpine experimental setup, IR mice were treated with CV four times per week for 60 days. Treatment was then halted at day 90, and physiological saliva secretion was measured bi-monthly (Figure 3A). As shown in Figure 3B and S3A, saliva output from IR+CV mice remained at non-IR control levels for 56 days after treatment termination (146 days post-IR). Unexpectedly, by 70-80 days post-treatment cessation (160-174 days post-IR), salivary flow declined to levels comparable to IR+S mice (Figure 3B, blue bars), indicating that the functional rescue of irradiated SGs gradually diminishes between 56 and 84 days (∼2-3 months) after treatment termination. This result questioned whether a minimum threshold level of endogenous muscarinic signaling is required for the regenerating tissue to hold a consistent and permanent homeostatic state.

**Figure 3.**
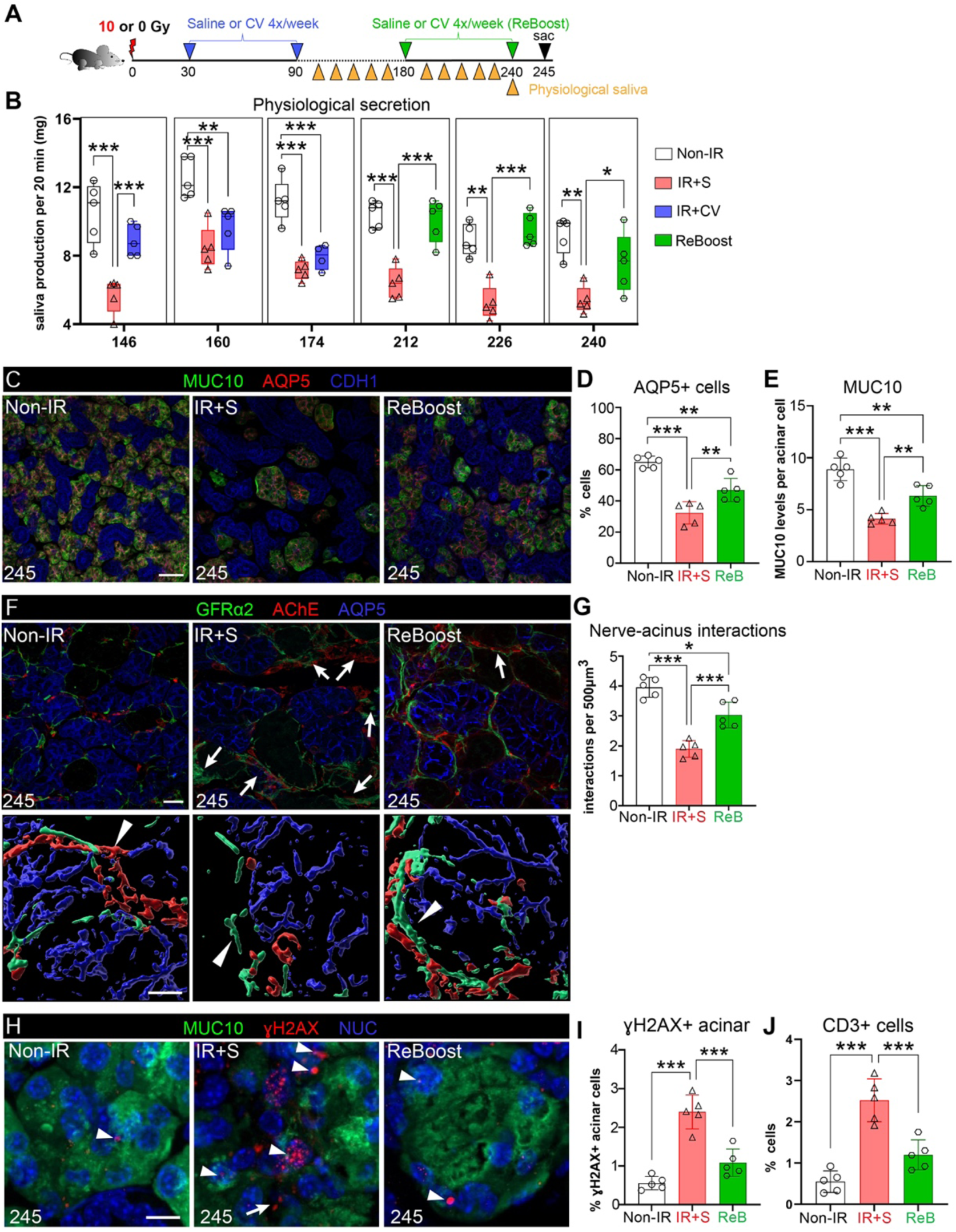
IR SGs remain highly responsive to an additional round of muscarinic treatment delivered months after radiation exposure. **A.** Schematic of the treatment regimen showing 1^st^ and 2^nd^ (i.e., ReBoost, ReB) application of saline (S) or muscarinic agonist cevimeline (12 mM) after IR. **B**. Physiological saliva secretion of the treatment groups measured from 146 to 240 days post-IR. **C-E**. Immunofluorescent staining and confocal imaging of AQP5^+^ and MUC10^+^ acinar cells in non-IR, IR+S, and IR+ReB SGs at 245 days post-IR (C), and quantification of representative acinar cell numbers and volume in SGs of each treatment group (D-E). **F-G.** Immunofluorescent analysis of GFRα2^+^ and AChE^+^ nerves surrounding acini (F), and quantification of nerve-acinar cell interactions (G) at 245 days post-IR across the 3 treatment groups. 3D surface-reconstructed images (F, lower panels) were generated using IMARIS. Arrows indicate presence of thick GFRα2^+^ and AChE^+^ nerves distant to acini, while arrowheads represent nerves interacting with acini. **H-I**. Immunostaining for MUC10 and γH2AX (H) at 245 days in the different groups (H). Arrowheads highlight γH2AX in acinar and arrows non-acinar cells. Quantification of γH2AX+ acinar cells from H (I). J. Quantification of CD3+ T cells in the different groups. Mean±SD. *, *P*<0.05. **, *P*<0.01. ***, *P*<0.001. Scale bars, 40 μm in C, and 20 µm (upper panel) and 10 μm (lower panel) in F and H.

**Figure S3.**
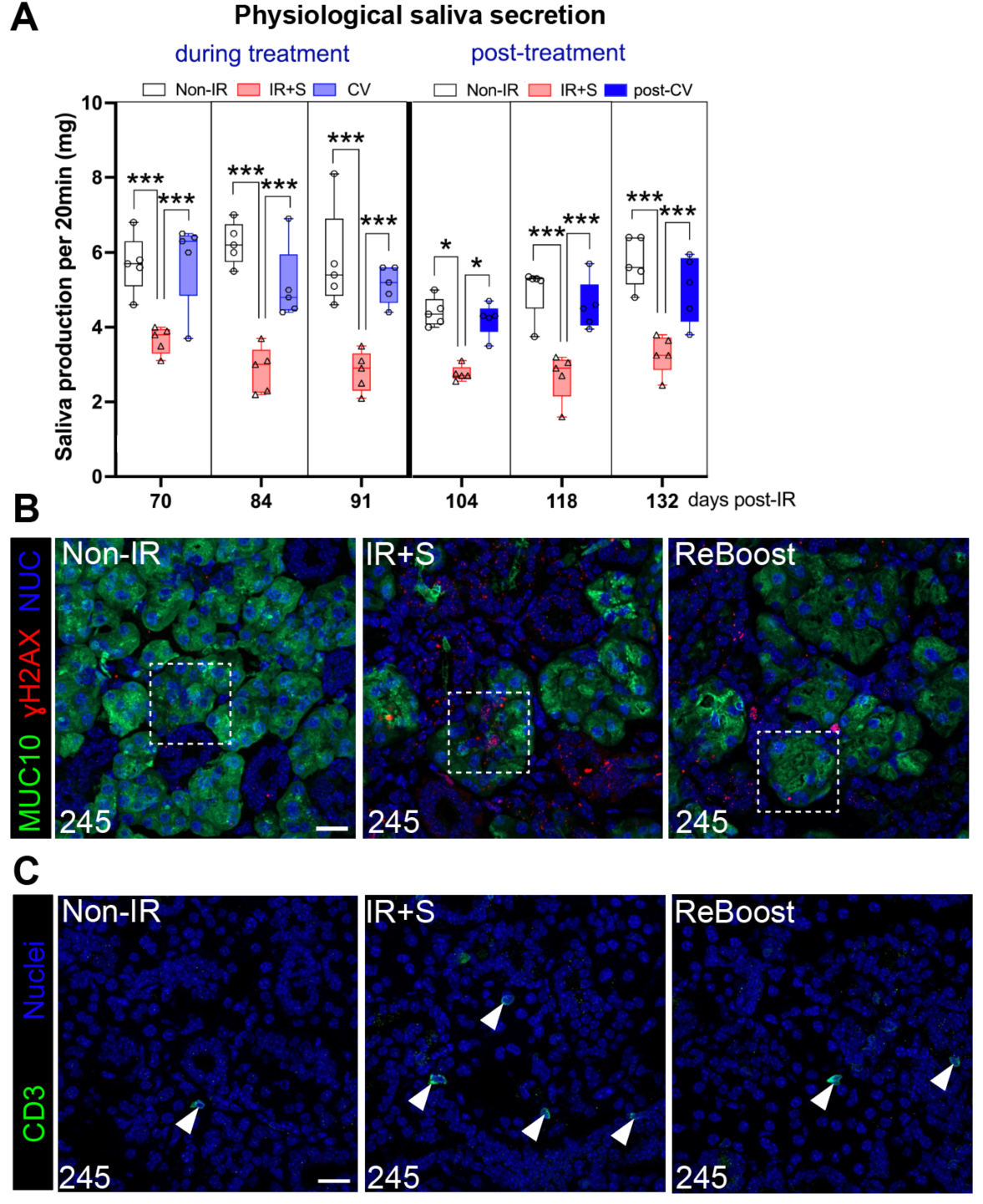
IR SGs remain highly responsive to an additional round of muscarinic treatment delivered months after radiation exposure, related to Figure 3. **A.** Physiological saliva secretion was measured during treatment (left) and after treatment termination at 90 days post-IR (right). **B.** Detection of DNA damage (γH2AX) foci in acinar cells across the 3 treatment groups (non-IR, IR+S, and IR+ReBoost with CV). Inset shown in Figure 5H. **C.** Immunostaining of CD3+ T cells (arrowheads) in SGs across the groups. Mean±SD. *, *P*<0.05. **, *P*<0.01. ***, *P*<0.001. Scale bars are 20μm.

Given this response, we questioned whether the suboptimal rescued IR+CV SGs remained receptive to CV stimulation and could regain the homeostatic state by providing a second round of external muscarinic stimulus (named ReBoost) at 180 days post-IR (6 months post-IR), 20 days after the functional endpoint (Figure 3A). Mice were treated with CV 4x per week for 60 days (180 to 240 days post-IR) and physiological saliva measured bi-monthly before mice were sacrificed 5 days after the last dose i.e., at 245 days or 8.5 months post-IR (Figure 3A, green). Remarkably, by 32 days of ReBoost saliva output had already returned to non-IR levels, contrasting to the continued low levels for mice re-treated with saline (Figure 3B). Furthermore, the increased saliva output was sustained at homeostatic-like levels up to the end of the testing period at 240 days post-IR (Figure 3B). As would be predicted for SGs showing improved salivary flow, acinar cells were greatly preserved at 245 days post-IR, albeit it at lower levels than non-IR SG, with the number of AQP5^+^ cells (Figure 3C-D) and MUC10 expression (Figure 3C-E) being significantly greater than saline-treated IR controls. Although nerve fibers in ReBoosted SGs were thicker and more frequently penetrated mesenchymal regions compared to non-IR controls (Figure 3F, arrows), nerve-acinar interactions were significantly enhanced relative to IR+S SGs (Figure 3G), supporting the observed improvement in physiological secretory function. Strikingly, the number of DNA-damaged acinar cells in ReBoosted SGs was comparable to non-IR tissue and markedly lower than in IR+S controls (Figure 3H-I, arrowheads, and Figure S3B). Together with the significant rescue of acinar cells, enhanced nerve-acinar interactions (Figure 3C-G), and reduced CD3+ T cell infiltration (Figure 3J and S3C) in ReBoosted SGs compared to the IR+S group, these findings suggest that the IR tissue continues to reside in a regenerated state when the minimally required muscarinic levels, provided solely endogenously and/or externally, are maintained in the tissue.

Thus, these data show that the function and structure of IR SGs are sustained for months after the end of muscarinic agonist treatment and that the rescued tissue retains a robust capacity to respond to regenerative cues, enabling the maintenance and restoration of tissue function.

### De-differentiated acinar cells are greatly expanded in chronically degenerating SGs, an outcome reversed by muscarinic agonist treatment

Acinar cells of the SG, along with those of the pancreas, can undergo de-differentiation in response to stress ^24–27^, an outcome that results in alterations in cell fate, such as acquisition of stemness^28^, ceasing of normal activity, and progression towards cellular dysfunction or cell death^29^. Based on these possibilities, we utilized single nucleus RNA sequencing (snRNA-seq) to investigate acinar cell identities and cell states in non-IR, IR+S, and IR+P (3 mM) SGs treated 30- to 90-days post-IR. Nuclei were isolated from each of the 3 groups (5 individual adult mice per group, 10 SGs pooled for each treatment) and subjected to 10x Chromium snRNA-seq. A total of 31,929 cells passed quality control procedures, with an average of 1,018 expressed genes per cell. Unsupervised clustering and differential expression analyses identified 9 major cell-type clusters (Figure 4A, left, and Figure S4A-B). Each cluster was annotated based on expression of recognized markers, including acini (*Aqp5*^+^), ducts (*Adcy2^+^*), myoepithelial (*Acta2^+^*), mesenchymal (*Dcn*^+^), endothelial (*Pecam1*^+^), nerves (*Ncam1*^+^), as well as immune (*Ptprc*^+^ and *Cd74*^+^), and pericyte (*Rgs5^+^*) cells (Figure S4A and B). Although our data showed the same cell type clusters regardless of condition (Figure S4C), the percentage of each cell type was altered between groups (Figure S4D). Compared to non-IR SGs, saline-treated tissue showed a substantial decrease in acinar cells (∼40%) and an increase in ductal cells (∼2-fold) as well as immune cells, including T cells (∼2-fold) and macrophages (∼2.9-fold) (Figure S4D), consistent with extensive tissue degeneration. As expected for a rescued tissue, the inverse was true for IR+P SG compared to IR+S (Figure S4D), supporting the restoration of cellular composition.

**Figure 4.**
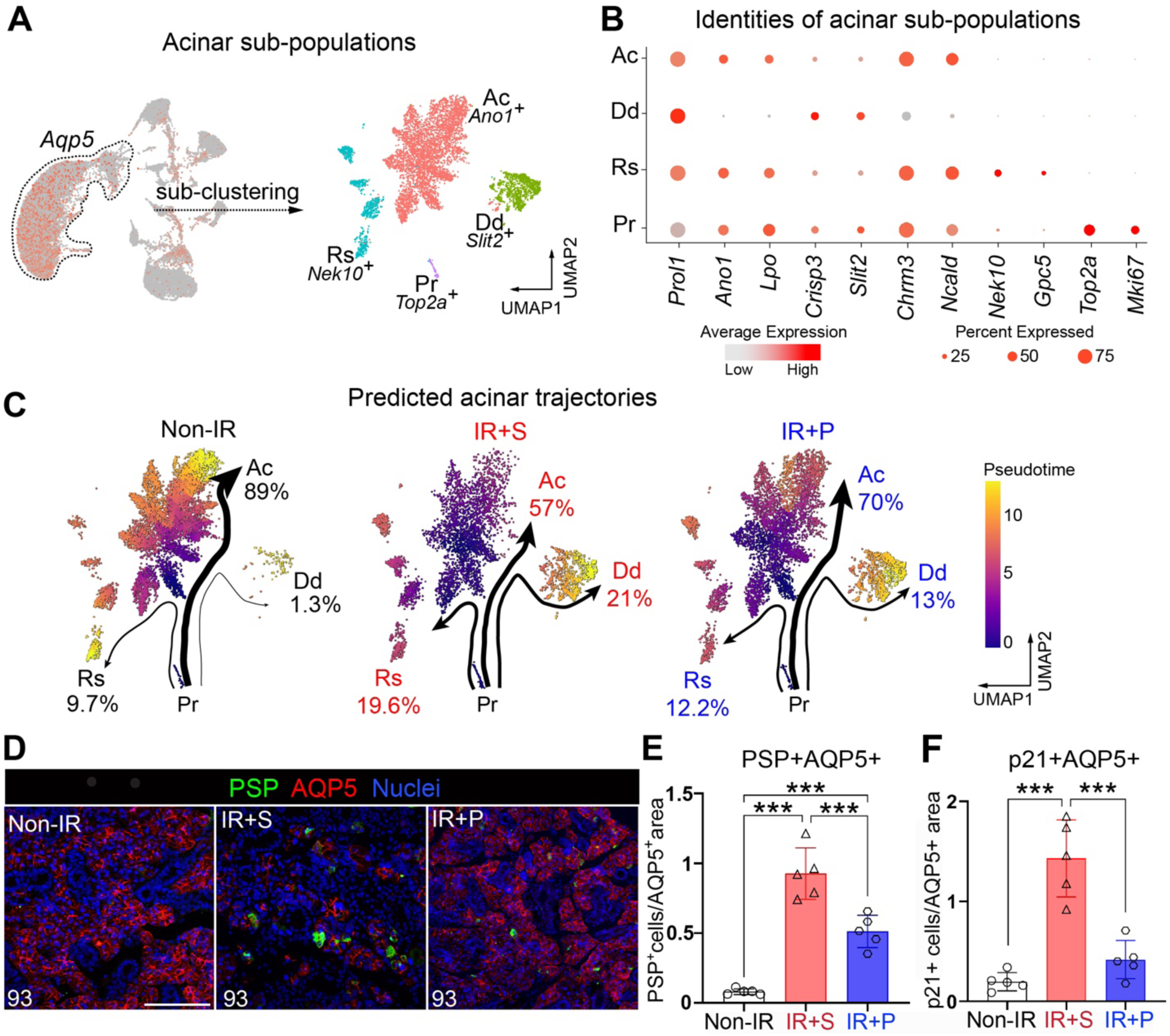
De-differentiated acinar cells are greatly expanded in chronically degenerating SGs, an outcome reversed by muscarinic agonism. **A**. Extraction and sub-clustering of the acinar cell snRNA-seq data identifies 4 distinct cell populations: active (Ac), de-differentiating (Dd), resting (Rs), and proliferating (Pr) acini. **B**. Dot blot showing enrichment in specific marker genes for each cluster, as seen in A. **C.** Predicted lineage trajectories for acinar clusters across the 3 different conditions. Pr cells were set as the pre-specified starting cluster. Thickness of arrows and percentages shown are associated with the number of cells in transition between clusters. **D-E.** Immunofluorescent analysis (D) and cellular quantification (E) of acinar cells expressing PSP (a marker of de-differentiation) in the different treatment groups at 93 days post-IR. Scale bar, 100 µm. Mean±SD, ***, *P*<0.001. **F.** Quantification of p21^+^ acinar cells across the different treatment groups. Mean±SD. ***, *P*<0.001.

**Figure S4.**
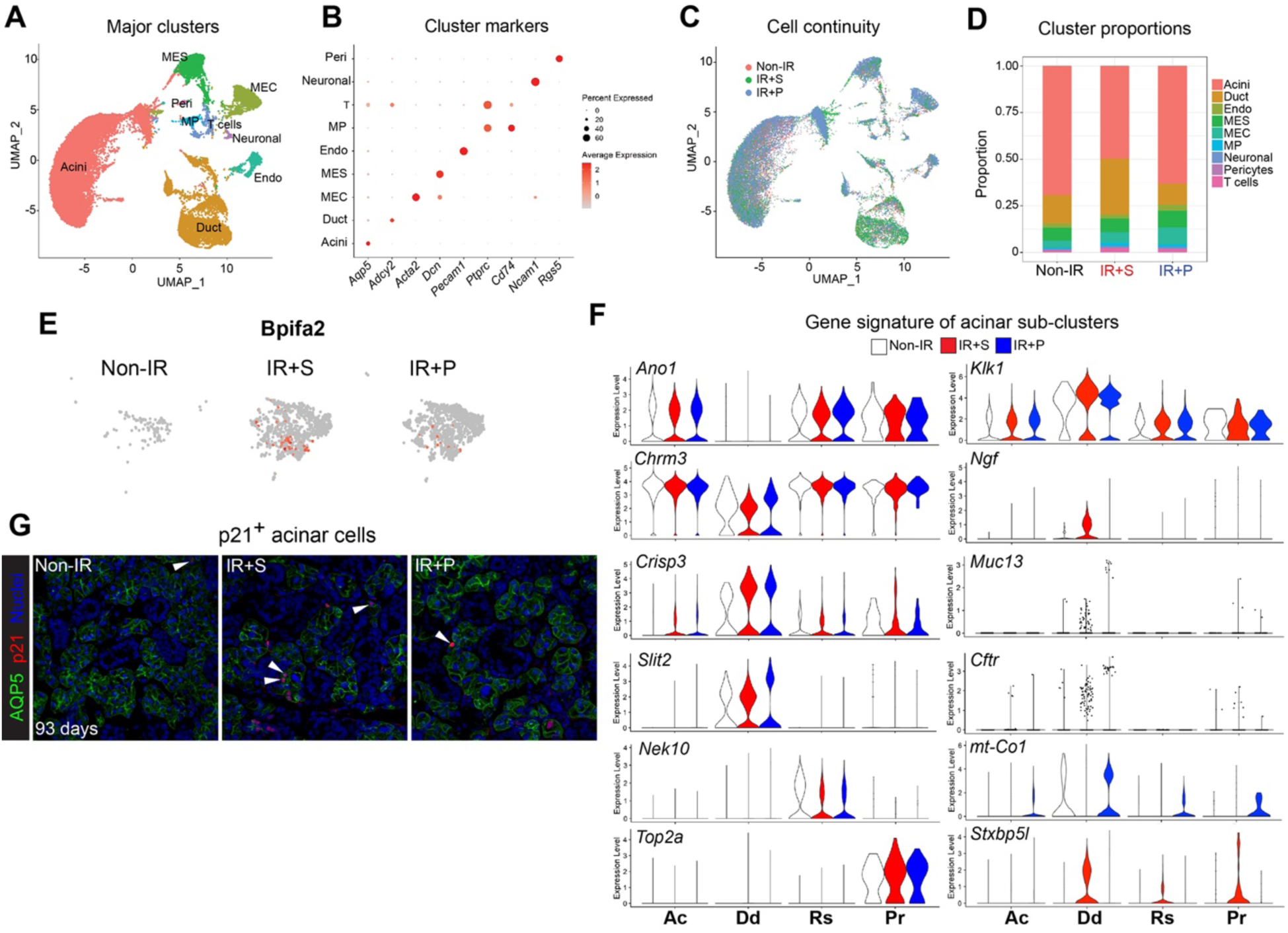
De-differentiated acinar cells are greatly expanded in chronically degenerating SGs, an outcome reversed by muscarinic agonism, related to Figure 4. **A.** Unbiased cluster analysis. MEC, myoepithelial cell. Peri, pericytes. MES, mesenchymal cell. Endo, endothelial. MP, macrophage. T, T cell. **B.** Integrated dot plot presents a specific marker gene for each of the 9 major clusters identified. **C.** UMAP plot of the integrated single nuclei from the 3 treatment groups, non-IR, IR+Saline (S), and IR+Pilocarpine (P, 3 mM) at 93 days post-IR. **D.** Bar plot showing the percentage of each cell type in each of the 3 treatment groups**. E.** *Bpifa2* is relatively enriched in de-differentiated (Dd) acini in the IR+S group, as compared to the non-IR and IR+P group. **F.** Violin plots showing gene expression signature in the 4 acinar sub-clusters across the 3 treatment groups. Ac, active. Dd, de-differentiated. RS, resting. Pr, proliferating. **G.** Immunostaining and confocal imaging of acini in cell cycle arrest, as marked by p21. Scale bar, 20µm. Arrows point to p21^+^AQP5^+^ acini.

Acinar cell identities were then determined through extracting cells based on expression of lineage specific genes, including *Aqp5, Prol1* (MUC10), and *Bhlha15* (MIST1)^30,31^. Sub-clustering of integrated samples identified 4 distinct populations within the acinar cell group for each condition (Figure 4A-B). One cluster was enriched in the calcium activated chloride channel *Ano1/Tmem16a* ^32^, the muscarinic receptor *Chrm3* ^33^, and functional saliva proteins essential for saliva secretion, thereby classifying it as active (Ac) acinar cells. A second acinar cluster was deficient in *Ano1* and *Chrm3* but highly enriched in *Bpifa2* (parotid secretory protein, PSP) (Figure 4A-B and S4E, Dd), a pro-acinar specific gene expressed during submandibular gland development and maturation but not by adult tissue^34^. This same cluster also showed expression of ductal-associated genes such as SG-related *Crisp3*^35,36^ and pancreas-related *Slit2*^37^ (Figure S4F), suggestive of acquisition of a de-differentiated phenotype in response to IR injury ^24–27^. Thus, this cluster was classified as a de-differentiated (Dd) acinar sub-cluster. The third cluster was exclusively enriched in cell cycle arrest genes, *Nek10* and *Gpc5*^38,39^, and was therefore termed as resting (Rs) acinar cells (Figure 4A-B and S4F). The fourth cluster distinctively expressed genes associated with cell division, such as *Top2a* and *Mki67*, thus marking them as proliferating (Pr) acinar cells (Figure 4A-B and S4F).

Based on these sub-clusters, we next analyzed potential differentiation trajectories of acinar subtypes in the 3 treatment groups using Slingshot^40^. Pr cells were chosen as the pre-specified starting cluster. Computed pseudotime analysis predicted 3 independent terminal trajectories moving from a proliferating to an active, resting or de-differentiating population, with the extent of cell state, shown by the thickness and length of the arrows (Figure 4C). In non-IR SGs, proliferating acini were predicted to majorly follow the path towards active acini (Ac: 89% of total cells), followed by resting acini (Rs: 9.7%) and a minor contingent of de-differentiated cells (Dd: 1.3%). In comparison, saline-treated IR SGs, drastically altered their predicted cells states, with a profound increase in proliferating cells moving towards Dd (from 2.3% to 21%) and Rs (from 9.7% to 19.6%) states, while the Ac sub-cluster was reduced (from 89% to 57%). Pilocarpine treatment strongly reversed these outcomes, resulting in a substantial reduction in the IR-induced generation of both Dd cells (21% to 13%) and Rs cells (19.6% to 12.2%) and an increase in Ac cells (from 57% to 70%; Figure 4C). In addition to the reduction in the number of Dd cells with pilocarpine, IR+P Dd cells also exhibited a reduction in the major muscarinic receptor *Chrm1* and ductal gene transcripts, including *Klk1* ^41^, *Ngf* ^42^, *Muc13* ^43^ and *Cftr* ^44^ compared to Dd cells in IR+S controls (Figure S4F), suggesting pilocarpine promotes the re-differentiation of this population into a more differentiated acinar-like cell. Moreover, in support of cells moving towards a more homeostatic or reparative cell state IR+P Dd cells were also enriched in genes associated with increased mitochondrial function (e.g., *mt-Co1*/COX1) (Figure 4SF), the terminal enzyme of the respiratory chain that is essential for aerobic energy generation^45^, whereas IR+S Dd cells were enriched in genes linked to senescence and fibrosis, such as *Stxbp5l* ^46,47^ (Figure 4SF). Thus, together these data point towards a role of muscarinic agonism in reestablishing functional, homeostatic acinar cells.

We further validated the reduction in Dd and Rs cells by immunostaining each group for the major de-differentiation marker *Bpifa2*/PSP and the cell cycle inhibitor CDKN1A/p21^48^. In support of our snRNAseq analyses, very few PSP^+^ acinar cells were present in non-IR SG (Figure 4D-E), consistent with the absence of injury, whereas saline-treated IR controls showed an 80-fold increase in these cells, with acini often composed of multiple PSP+ cells (Figure 4D-E). Pilocarpine treated IR SG, however, showed an almost 50% reduction in the PSP^+^ population compared to saline-treated controls, suggesting that the acinar population resembled that seen in the homeostatic state (Figure 4D-E). This regenerative outcome was also strongly supported by the alteration in p21+ acinar cells. In line with the trajectory analysis revealing IR-induced injury to promote a more resting state, IR+S SGs exhibited a 3.5-fold increase in the number of acinar cells expressing p21 as compared to non-IR SG (Figure 4F and S4G). Pilocarpine-treatment significantly reversed this outcome, with the number of p21^+^ cells being similar to non-IR SGs (Figure 4F and S4G), thus revealing muscarinic agonism to provoke cell cycle re-entry required for eliciting proper acinar cell replenishment.

In summary, these data suggest radiation-induced tissue degeneration is not only driven by acinar cell loss but also by the transition of IR-surviving acinar cells towards a de-differentiated, growth arrested cell state and that this can be greatly reversed through muscarinic activation.

### Muscarinic-receptor induced calcium signaling reestablishes mitochondrial metabolism and differentiated acinar cells

Secretory function encompasses the continuous production of secreted proteins and cell contraction, and as such, requires a constant energy supply. A plethora of studies in other tissues show that mitochondrial dysfunction impairs tissue regeneration^49–51^, implying that IR SG degeneration is due, in part, to disrupted mitochondrial metabolism. Consistent with this, analysis of RNAseq datasets at 30 days post-IR showed a critical downregulation of mitochondrial metabolism-associated pathways, including “oxidative phosphorylation” and “ATP synthesis” and related genes (e.g., *ATP5/Atp5* family members) (Figure 5A and S5A). Similarly, the numbers of active mitochondria (marked by MitoTracker^+^) and levels of adenosine triphosphate (ATP, BioTracker^+^) in acinar cells were greatly diminished compared to non-IR SGs (Figure 5B and S5B), consistent with the non-healing state of the tissue. This outcome was also apparent in human IR SGs (Figure S5C-D, publicly available RNAseq datasets), indicating a conserved correlation between mitochondrial dysfunction and SG degeneration.

**Figure 5.**
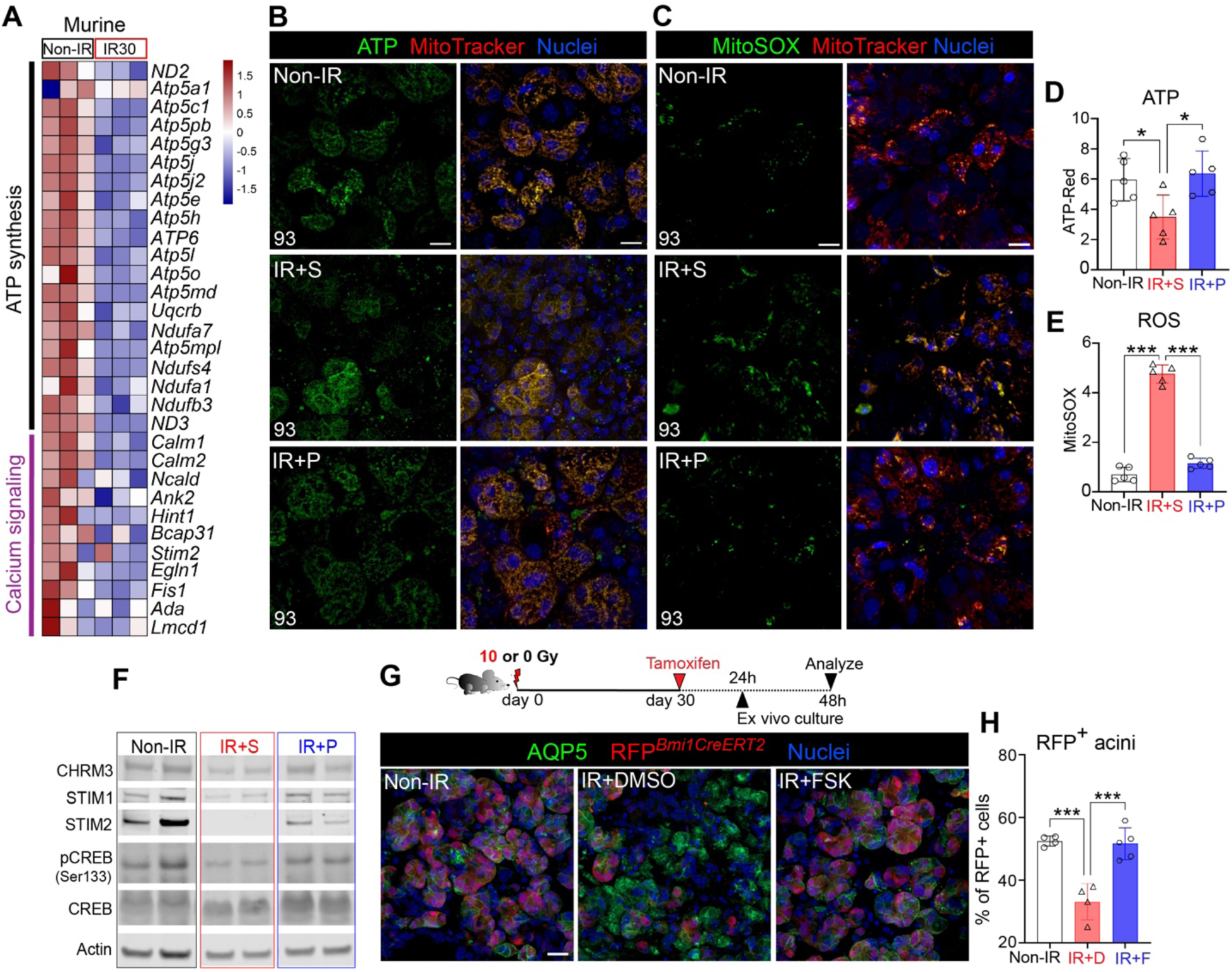
Muscarinic-receptor induced calcium signaling reestablishes mitochondrial metabolism and differentiated acinar cells. **A.** Heatmap of ATP synthase and calcium signaling-related gene expression levels at 30 days post-IR and control (Non-IR) murine SGs. **B-C.** Live imaging of ATP (D) and ROS (MitoSOX) (E) in non-IR, IR+S, and IR+P (3 mM) SGs at 93 days post-IR. MitoTracker, active mitochondria. **D-E.** Quantification of ATP and ROS in B and EC, respectively. **F.** Western blot analysis of proteins associated with calcium signaling across the 3 different treatment groups. Protein levels were normalized to actin. **G-H.** Schematic shows the experimental setup for ex vivo lineage tracing of *Bmi1^CreERT^*^2^*;Rosa26^RFP^* SG rudiments. Mice were irradiated with 0 or 10 Gy and lineage tracing induced at 30 days post-IR. SGs were extracted 24 hrs later and cultured for 24 hrs in DMSO (IR+DMSO, labelled IR+D in J) or forskolin (FSK, Ca^2+^ activator, IR+F in J). non-IR SG rudiments were treated with DMSO. Tissue was immunostained for AQP5. The percentages of RFP^+^ cells in I are quantified in H. Mean±SD. *, *P*<0.05. **, *P*<0.01. ***, *P*<0.001. Scale bars in in B, C, and G are 20 µm.

**Figure S5.**
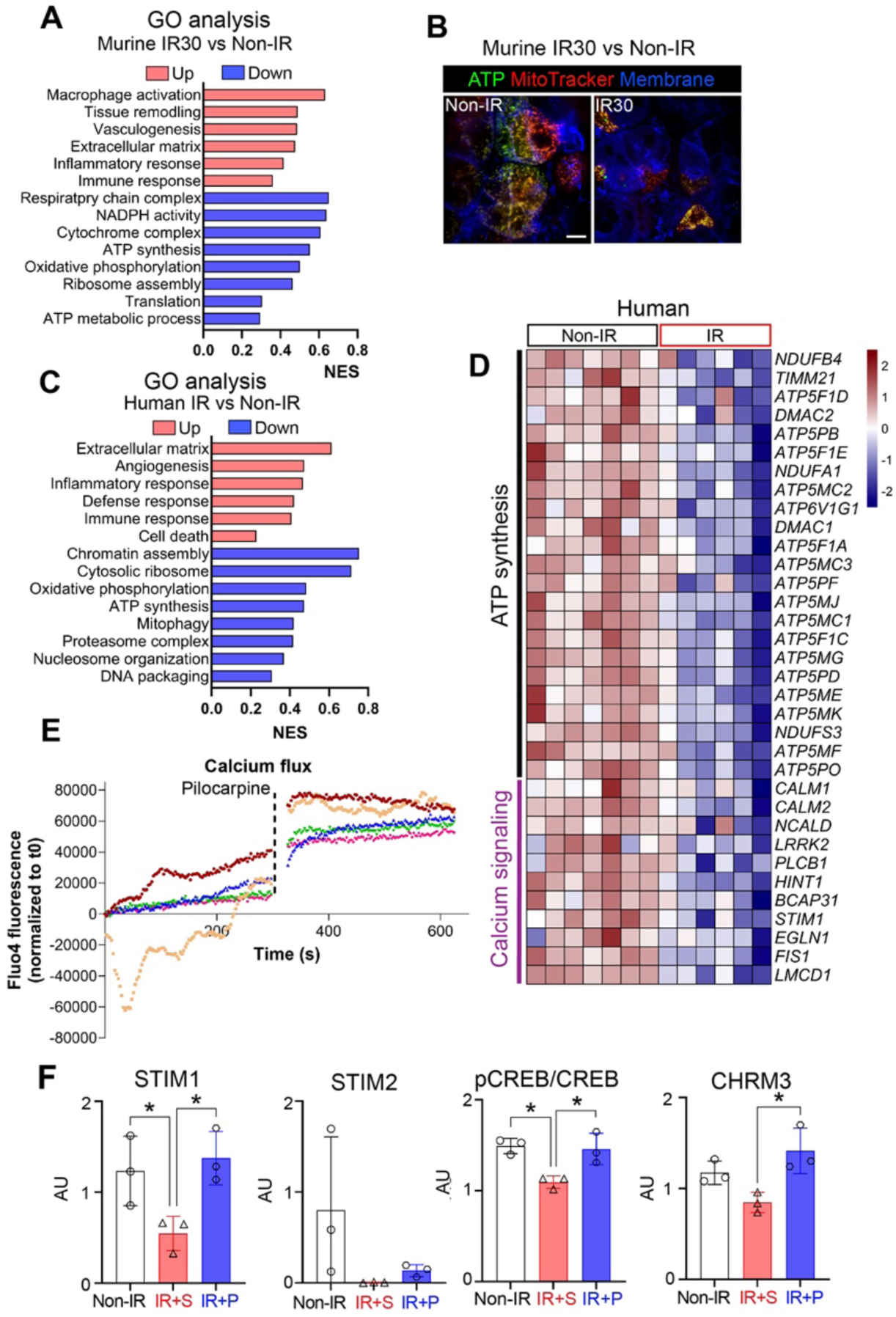
Muscarinic-receptor induced calcium signaling reestablishes mitochondrial metabolism and differentiated acinar cells, related to Figure 5. **A.** Gene Ontology (GO) analyses of murine non-IR versus 30 days post-IR SGs bulk RNA-seq datasets. **B.** Live imaging of ATP synthesis and active mitochondria (MitoTracker) in non-IR and 30 days post-IR murine SGs, cultured ex vivo for 24 hours. **C.** GO analysis of human non-IR versus IR SGs (>2 yrs post-radiation, GSE206878^26^) bulk RNA-seq datasets. **D.** Heatmap of ATP synthase and calcium signaling-related gene expression levels in human salivary glands, irradiated (IR) versus healthy (Non-IR). **E.** Calcium signaling levels, with or without muscarinic stimulation, measured by live imaging of Fluo-4 AM uptake by freshly harvested SGs. Each color represents a biological replicate **F.** Quantification of STIM1, STIM2, pCREB and CHRM3 protein levels in each condition determined by western blot (Figure 5F). Groups were normalized to actin. Mean±SD. *, *P*<0.05.

Acute injury results in a transient increase in ROS production, where it aids in the repair process, and decrease in ATP that return to homeostatic levels upon the completion of wound repair^52^. However, an overabundance of ROS in response to severe chronic injury, such as ionizing radiation^52^, results in chronic oxidative stress and subsequently, delayed wound healing and chromosomal instability ^53,54^. Although treatment of different cell types pre-injury has shown muscarinic agonism to protect mitochondrial function, reduce production of ROS or prevent DNA damage DNA damage^55^, it is unknown whether degenerated tissues can respond in such a fashion. To test this possibility, mice were treated from 30-90 days with pilocarpine and SG explants at 93 days post-IR were immediately assayed for changes in mitochondrial function through live imaging. Consistent with an upregulation of oxidative stress during radiation-induced degeneration, ROS was dramatically increased by 400% in saline-treated SGs compared to non-IR SGs at 93 days post-IR (Figure 5C and E; ROS marked by MitoSOX) whereas ATP was significantly decreased by 40% (Figure 5B and D; MitoTracker). In complete contrast, redox reactions were essentially restored with pilocarpine-treatment (Figure 5B-E), with levels of ROS and ATP mimicking those of non-IR controls (Figure 5B-E), indicating that muscarinic activation is able to dampen chronic oxidative stress, thereby allowing tissue to re-enter the wound healing program.

Next, we questioned the mechanism through which muscarinic agonist reverses mitochondrial dysfunction and promotes acinar restoration. A key regulator of mitochondrial metabolism in repair and regeneration is intracellular calcium release, which is also activated by muscarinic stimulation ^56,57,58^. Transcriptomic analysis of IR versus non-IR SGs at 30 days post-IR revealed calcium-related gene transcripts (e.g., *STIM2/Stim2*, *NCALD/Ncald*) in IR mice (Figure 5A) and human SG to be significantly downregulated as compared to non-IR tissue (Figure S5D), indicating a downregulation of calcium-regulated events in the degenerated SG. Given this response, we asked whether re-activation of Ca^2+^ signaling through muscarinic agonism (Figure S5E) could rescue redox homeostasis in IR SG. Analysis of murine STIM1 and 2 protein - critical sensors of endoplasmic reticulum (ER)-luminal Ca^2+^ levels that maintain a cellular Ca^2+^ balance - in day 90 post-IR SG extracts revealed both proteins to be significantly downregulated or absent in IR+S SGs, as compared to non-IR SGs (Figure 5F and S5F). Similarly, phosphorylation of CREB (pCREB), a readout of increased intracellular Ca^2+^ ^59,60^, was also significantly downregulated (Figure 5F and S5F). In contrast, pilocarpine treatment (IR+P 3mM) effectively reversed these outcomes, as shown by the significantly increased levels of STIM2 (regulates store-operated and store-independent Ca2+-influx) and pCREB (calcium-responsive transcription factor), as well as an increasing trend of STIM1, as compared to the IR+S group (Figure 5F). In addition, CHRM3 expression was also significantly increased in IR+P SGs, as compared to IR+S group (Figure S5F).

Given these outcomes, we questioned whether Ca^2+^ activation by muscarinic agonism was directly linked to the replenishment of acinar cells in degenerating SG through genetic lineage tracing in an ex vivo setting. To test this hypothesis, we tested acinar cell replacement through genetic lineage tracing using tamoxifen-inducible *Bmi1^CreERT^*;*Rosa26^RFP^* mice^22^. *Bmi1^CreERT^*;*Rosa26^RFP^* mice were irradiated with 0 or 10 Gy, and lineage tracing induced at 30 days post-IR by tamoxifen injection. Twenty-four hours later, SG tissue was harvested and cultured ex vivo for a further 48 hours in the presence of DMSO or forskolin (FSK), a cell permeable adenylate cyclase activator that induces calcium signaling^61^ (Figure 5G, schematic). Quantification of lineage-traced cells revealed a 56% decrease in RFP^+^ acinar cells in DMSO-treated IR SG compared to healthy non-IR controls (Figure 5G-H). In contrast, culture of IR tissue with FSK led to near global repopulation of acinar cells, with the percentage of RFP^+^ cells matching that of non-IR controls (Figure 5G-H), thus confirming a direct influence of Ca^2+^ in acinar cell replenishment during the repair of degenerated SGs.

Altogether, these data strongly support the stimulation of calcium-mitochondrial oxidative metabolism as a rescue mechanism to replenish acinar cells, and likely other cell types, in chronically degenerating SG (Figure 6).

**Figure 6.**
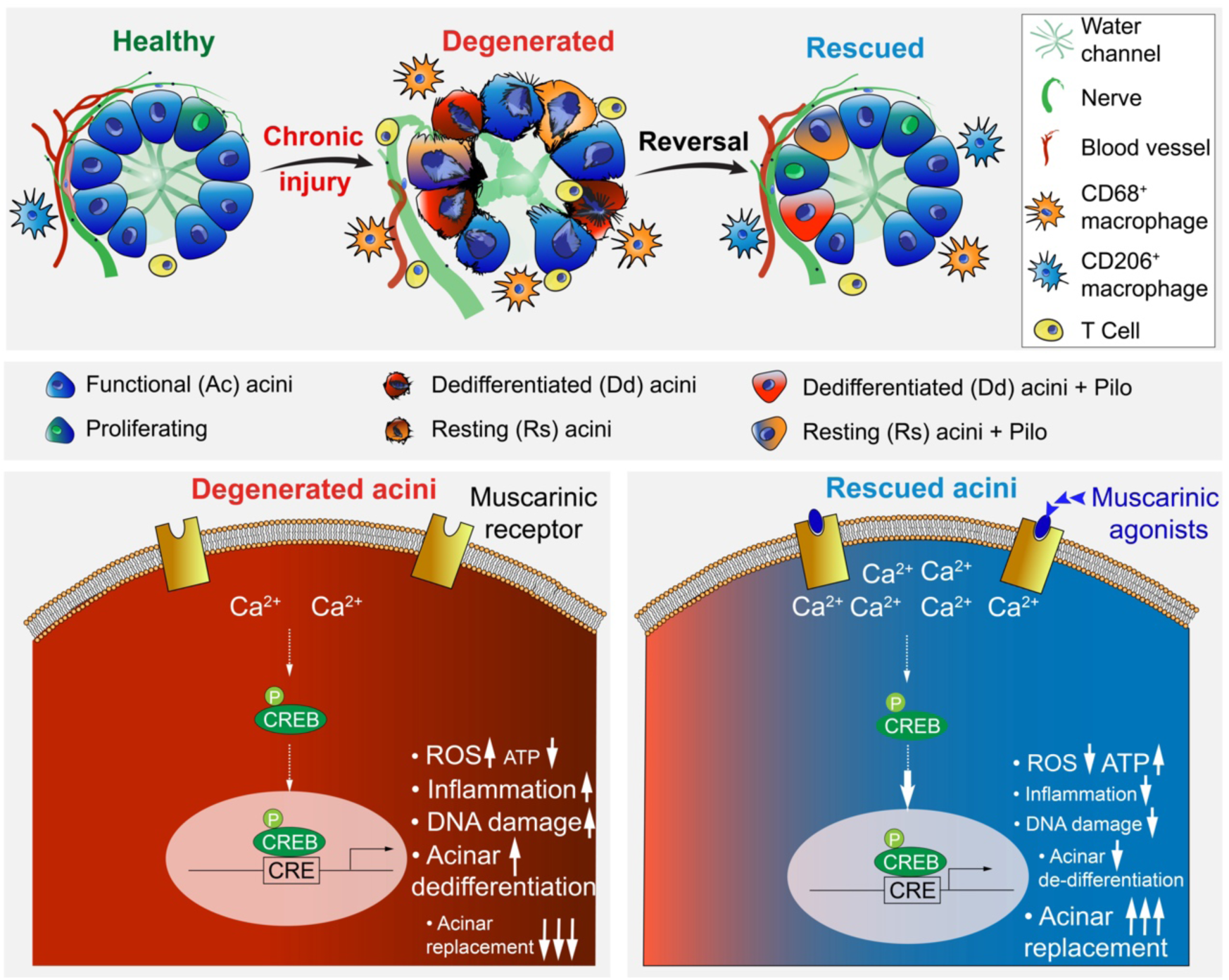
Chronic degenerating salivary glands can be rescued by restoring mitochondrial function.

## DISCUSSION

The continual absence of regenerative solutions for chronically damaged non-healing organs has implied that progressive degeneration causing organ failure is irreversible. For tissues of the head and neck area, as well as lung, heart, skin and abdomen/pelvis region, chronic radiation injury causes long-term complications that are clinically treated by a variety of symptomatic tools, such as anti-inflammatory agents, hyperbaric oxygen, topical analgesics, and steroids^63^. In addition to similar symptomatic agents, patients can also be treated for damage prevention and tissue preservation when given in early phases post-damage. For example, in the SG a number of potential radioprotective therapies, such as P2X7R antagonists^64^, KGF^65^, and amifostine^66^, have shown positive pre-clinical/clinical results in reducing damage, however, translation and integration into clinical practice has been relatively poor for various reasons ^67,68^. Similar to other organs^69–71^, regenerative treatments are mostly aimed at activating SG repair during acute time points - including AAV2-sonic hedgehog (Shh^72^), ABT-263^73^, CERE-120^74^, and GDNF^75^. These are promising approaches, but possess little to no efficacy when tested at chronic stages of tissue degeneration in small and large animal models^72,74,76^. Additional therapies aimed to regenerate chronically damaged organs, which are based on transplantation of epithelial^74^ and mesenchymal stem cells^67,68^ or gene therapy^74,75^, progressed to clinical trials and are currently being tested. However, no data is yet available that suggests that these therapies are effective in fully rescuing a chronically degenerated gland long-term in treated patients. Not disregarding the potential of these therapies to be reparative, our findings show a strong tissue wide and sustained reversal of chronically injured SGs after neuromimetic treatment. Based on nerve-derived signals being essential for maintaining many other organ/tissues, we predict that other epithelial organs adversely impacted by chronic injury may respond in a similar restorative fashion, thus opening up the clinical probability of repairing various degenerative conditions.

Many regenerative processes have been associated with cells acquiring an alternate state dedifferentiation^78,79^. Substantial work has been performed in the pancreas showing terminally differentiated acinar cells revert back to a less-differentiated state in response to injury (e.g., pancreatitis). During this process there is a shift from an acinar cell to a dedifferentiated, ductal-like phenotype, that correlates with reduced expression of acinar genes and an increase in ductal and progenitor-lie markers markers^29^. These dedifferentiated cells subsequently act to repair and regenerate the tissue, at least in the event that there is no further perturbation that leads to sustained chronic degeneration such as fibrosis^80^. In support of this general concept of acinar plasticity in response to damage, we also identified a novel dedifferentiated acinar cell type (Dd) that specifically increases after radiation injury. Due to the severity of the damage and the global loss in functional acinar cell number, this Dd cell population is presumably not able to heal the gland. However, in addition to muscarinic treatment reducing the number of Dd cells, the molecular status of these Dd cells also changed, with acquisition of a more homeostatic-like acinar state. Whether these alterations actively enhance tissue regeneration or organ secretion remains unclear and requires further investigation.

Since mitochondrial dysfunction and oxidative stress are hallmarks of many other degenerative conditions^81^, it is not surprising that mitochondrial dysfunction, or the inability to respond to high energy requirements under stress, has already been associated with acute radiation injury in SGs^82–84^. During this acute phase, elevated ATP^82^ and ROS^83^ in mitochondria of acinar cells activates a TRPM1-mediated increase in intracellular calcium ([Ca^2+^]_i_) that lowers fluid secretion, reduces ATP5, COX, and NAD^84^, and increases cell death. Our analysis of chronic injury highlights drastic alterations in cellular metabolism at the late degenerative stages, with upregulation of mitochondrial dysfunction and cell exhaustion, potent drivers of chronic disease and tissue destruction. The success of muscarinic stimulation in returning mitochondrial dynamics to homeostatic-like levels, in part, through re-balancing Ca^2+^-ATP dependent metabolic signaling, and the expression of muscarinic receptors 1 and 3 by a plethora of epithelial tissues, suggests that targeting this pathway may offer a regenerative opportunity for other chronically damaged epithelial organ systems. In the future it will be important to identify the genes lying downstream of this mitochondrial pathway in both the degenerative phase and upon muscarinic treatment to reveal the primary drivers of the regenerative process.

Radiation at levels typically delivered for the elimination of tumors predominately induces an irreparable state due directly to DNA damage and indirectly through the toxic production of free radicals such as ROS damaging DNA, lipid and protein damage^85^. As for other tissues, radiation-induced DNA damage in the SG results in apoptosis and senescence and has led to the investigation of agents that prevent damage or promote DNA repair. For example, murine studies using pre-IR delivery of agents that impair cell cycle entry (performed to increase the time for DNA repair) such as IGF1^86^ and the cyclin-dependent kinase inhibitor Roscovitine ^87^ or agents that prevent cell senescence (e.g., IL-6^20^) show a reduction in DNA damage and a benefit to SG function. The ability of muscarinic agonism to achieve extensive and sustained DNA repair in the chronically injured SG that is achieved without impairing cell cycle progression or preventing cell senescence points towards a new mechanism through which DNA repair can be effectively promoted. Further studies are needed to determine whether this novel outcome is also mediated via reactivation of mitochondrial metabolism or an alternative mechanism.

In summary, we demonstrate that degeneration of chronically injured organs can be reversed and that the functional and structural rescue is sustained long term, independent of treatment. Thus, we provide a novel therapeutic paradigm for organ restoration that may significantly benefit a diversity of chronic disease conditions.

## STAR METHODS

### Animal handling

All animals used in the study were housed in the Association for Assessment and Accreditation of Laboratory Care (AAALAC)-accredited University of California, San Francisco (UCSF) Laboratory Animal Resource Center. All procedures were approved by the UCSF Institutional Animal Care and Use Committee (IACUC), adhered to the NIH Guide for the Care and Use of Laboratory Animals, and executed under an approved IACUC protocol. The following mouse strains were obtained from the Jackson Laboratory (Bar Harbor, ME): C57BL/6J (JAX000664), *Bmi1-CreERT* (MGI: 3805814^22^) and *Rosa26-RFP* (MGI: 3809524^87^).

### γ-radiation of salivary glands

Female C57BL/6J or *Bmi1*-*CreERT*; *Rosa26-RFP* (age 6- to 7-weeks) mice received a single 10Gy dose of γ-radiation 30, 60 or 90 days before treatments as previously described^8,12^. Briefly, animals were anesthetized using 2.5% 2,2,2-Tribromoethanol (Thermo Fisher Scientific, A18706.22) and placed into the Shepherd Mark I Cesium Irradiator (JL Shepherd & Associates), with the body and cranial regions shielded from radiation using lead blocks. The lower head and neck regions of mice were exposed to radiation at a dose rate of 167 rad/min for 6 min for a total dose of 10Gy. A control group of mice were also anesthetized under the same conditions but did not undergo radiation treatment. Well-being of the irradiated mice was monitored for the first 48 hours. To ensure adequate food intake, all mice were given a soft diet *ad libitum* (clear H_2_O). Mice were then euthanized at 14-, 30-, 60-, 93-, 123-, or 245-days after radiation.

### Muscarinic agonist treatments

To prepare stock solution, 1.7mg of pilocarpine hydrochloride (Sigma-Aldrich, P0472) powder was dissolved in 25μL of sterile saline solution (Aspen, AHI14208186) to make a concentration of 300mM, and then filtered with a 0.22μm filter. A 1:100, 1:50 or 1:33.3 dilution was made before injection to achieve a final concentration of 3, 6 or 9 mM. Pilocarpine was administered by i.p. at 200μL/30g body weight with the designated concentrations 4 times per week for the duration of 2 months. Saline was also administrated accordingly as vehicle control. Cevimeline (Sigma-Aldrich, SLM007) at 12mM was injected by i.p. at 10 mg/kg body weight 4 times per week from 30 to 90 and 180 to 240 days post-IR.

### Physiological saliva collection

Citrate-induced gustatory stimulation (mornings at 9:00 a.m.) was utilized, as previously published with modifications^8^. Briefly, an absorbent filter paper (Bio-Rad, 1703932) was incubated in sodium citrate (73.6 mg/mL; Spectrum Chemical, S1250) solution for 15 min at room temperature (RT) on an orbital shaker and then dried overnight at RT. The filter paper was then cut into 30 mm × 2 mm strips, and the middle point was labeled on each strip. Individual strips were then placed in 1.5 mL microcentrifuge tubes and weighed. After mice were anesthetized by 2% inhaled isoflurane, a strip of filter paper was inserted into the oral cavity on top of the tongue until the maxillary incisors reached the middle point of the strip. After 20 min, the strip was removed and placed in the original 1.5 mL microcentrifuge tube. The amount of saliva collected was determined by measuring the difference in weight of the microcentrifuge tube with filter paper before and after collection, using a precision scale (OHAUS Adventurer).

### Immunofluorescent analysis

For immunofluorescent analysis, submandibular tissues were manually dissected and immediately fixed with 4% paraformaldehyde (PFA) at 4°C overnight, while skin samples were fixed for 15 min^88^. Fixed samples were thoroughly washed with 1x phosphate-buffered saline (PBS), cryoprotected by immersion in 12.5 and 25% sucrose solution, then embedded in optimal cutting temperature compound (O.C.T., Sakura Finetek USA, 4583) and stored at −80°C. Tissues were sectioned (12 μm or 20 μm) with a cryostat (Thermo Fisher Scientific) for immunofluorescence staining, as described below. Sectioned slides were permeabilized with 0.5% Triton X-100 (Thermo Fisher Scientific, A16046.AP) in PBS for 15 min at room temperature (RT), followed by 2 hours’ blocking at RT with 10% donkey serum (Jackson ImmunoResearch, 017-000-121), 5% BSA (Fisher BioReagents, BP9700100), and Mouse on Mouse (M.O.M) immunoglobulin G-blocking reagent (Vector Laboratories, BMK-2202) if required in 0.01% PBS-Tween 20. Then tissue sections were incubated with the corresponding antibodies (details in Table S1) overnight at 4°C. Antibodies were detected using Cy2-, Cy3-, or Cy5-conjugated secondary Fab fragment antibodies (1:300; The Jackson Laboratory), and nuclei were stained using Hoechst 33342 (1:1000; AnaSpec Inc., AS-83218). Slides were mounted using Fluoromount-G (SouthernBiotech, 0100-01).

### Quantification of immunofluorescent images and specific cell populations

For each treatment group, tissues were sectioned (via cryostat) from the exterior to the interior of the gland. Every second tissue section was taken for immunostaining, with each section being *x* = 3.5 mm, *y* = 3 mm, and *z* = 12 μm in size. Images were captured on an inverted Zeiss LSM 900 confocal microscope (RRID:SCR 022263) using a 10, 40 or 63X oil lens, depending on the application. Three to five fields of view for each tissue section were imaged with the size of each image measuring *x* = 250 μm, *y* = 250 μm, and *z* = 12 μm. The *z* axis was composed of 1-μm sections consolidated into a 12-μm projection. Fields of view were selected at three distinct locations across the tissue section (0.5 to 1 mm apart) and based on enrichment in acinar cells. For high magnification acquisitions, images were acquired with pinhole set to approximately 1μm and a 1μm step size was used for Z stack collection. Maximum intensity projections of sub-stacks specifically containing each layer were made using ZEN software (Zeiss, RRID:SCR 013672).

Acinar cells, myoepithelial cells, endothelial cells, and melanocytes were identified on the basis of AQP5 or MUC10, SMA, CD31 and DCT, respectively. Nerves were identified on the basis of TUBB3 (pan-neuronal), GFRα2 (parasympathetic) or TH (sympathetic) expression, respectively. Proliferating and DNA damage cells were identified on the basis of γH2AX expression, respectively. Immune cells were identified on the basis of the T cell marker CD3. Quantification of cells was achieved by applying the image processing software IMARIS. The number of acinar cells was normalized to the total number of nuclei per section. The number of DNA damaged cells was normalized to the total number of acinar cells. The number of immune cells was normalized to the total number of nuclei per section. Each quantification was performed on three to five sections per gland, from at least four animals.

### 3D image generation via IMARIS processing

Quantification of GFRa2^+^AChE^+^parasympathetic nerves was achieved through analysis of 256 μm × 256 μm images within a 12 μm × 1 μm projection. Three to six separate regions of each gland that were enriched in acinar cells were imaged per animal (5 animals per group). Images were then analyzed with IMARIS v9.6 (Bitplane), which provides unbiased measurements of immunofluorescence. Processed images were subjected to Gaussian filtering and background subtraction. Surface reconstructions were made with the “Surfaces” module. Object-based colocalization analyses was then performed using the “shortest distance” filter to quantify touching objects. Co-localized volumes were recorded for each of the groups (n=5 per group), and then visualized with Prism GraphPad as described in the “Statistical Analysis” section.

### Bulk RNA sequencing of salivary glands

For RNA analysis, tissue was snap-frozen and stored at −80°C. Total RNA was isolated from four 50 μm tissue sections of non-IR or 14-, 30-, 60- and 90-day post-IR submandibular glands (n=3) using the RNAqueous Micro Total RNA Isolation Kit (Thermo Fisher Scientific, AM1931), and treated with DNAse I (Thermo Fisher Scientific, AM1931) to remove genomic DNA. RNA integrity number (RIN) and yield was verified using the RNA Nano 6000 Kit (Agilent Technologies, 5067-1511) and a Bioanalyzer 2100 instrument (Agilent Technologies, RRID:SCR 019715). RNA sequencing of samples with RIN>7 was performed by Novogene (https://en.novogene.com) using the Illumina NovaSeq 6000 platform. Raw data (FASTQ) were processed through fastp^89^ to remove reads containing adapter and poly-N sequences and reads with low quality from raw data. Paired-end clean reads were aligned to the *Mus musculus* mm10 reference genome using the Spliced Transcripts Alignment to a Reference (STAR) software^90^. FeatureCounts was used to count the read numbers mapped to each gene^91^. Differential expression analysis between two conditions was performed using DESeq2 R package^92^. Genes with an adjusted P value of <0.05 were assigned as differentially expressed. Heatmap of differentially expressed gene clusters were visualized using pheatmap^94^. GO of differentially expressed genes was performed using g:Profiler^95^. Normalized Enrichment Scores (NES) were plotted using Prism 10 (GraphPad).

### Single-nucleus RNA sequencing

Single nuclei were isolated with Singulator S100 (S2 Genomics) from fresh frozen submandibular glands of non-IR, 93 days post-IR with saline and 93 days post-IR with pilocarpine treatments. The machine was set at nucleus mode with standard protocol. The isolated nuclei were then dyed with DAPI at 1:1000 and purified by flow cytometry. 40,000 of single nuclei from each group were submitted to UCSF Genomics Core and the UCSF Institute for Human Genetics Genomics Core, for library preparation and sequencing, respectively. Single nuclei were processed on the Chromium Controller (10x Genomics) using the Chromium Next GEM Single Cell 3’ Reagent Kits v3.1 (Dual Index) following the manufacturer’s instructions. Sample quality controls and quantification was assessed using TapeStation (Agilent Technologies), then sample libraries were sequenced on NovaSeq6000 platform (Illumina) generating 450M read counts for each GEM well. Raw FASTQ files were processed via CellRanger (v7.0.1, RRID:SCR_017344) on 10x Genomics Cloud Analysis (https://www.10xgenomics.com/products/cloud-analysis) to map to the reference genome (GRCm38) and generate filtered gene-barcode matrices.

R scripts were obtained from previous studies^34^. Briefly, filtered gene-barcode matrices were analyzed using the Seurat^95^ v4.097. Seurat objects were generated with CreateSeuratObject (min.cells = 3, min.features = 200). Cells were filtered based on the distribution of number of genes (nFeature, >200 and <5000) and percent mitochondrial genes (percent.mt <5) per cell. Data were normalized for sequencing depth, log-transformed, and multiplied by a scale factor of 10000 using the default parameters of NormalizeData. The top 2000 variable genes within each dataset were selected based on a variance stabilizing transformation and used in downstream principal component analysis (PCA). Clustering was performed at resolutions calculated by CellFindR^97^. Cell clusters were identified by construction of a shared nearest neighbor graph (FindNeighbors) and a modularity optimization-based clustering algorithm (FindClusters) using the PCs determined by PCA (dims = 1:20). Cells and clustering were visualized using Uniform Manifold Approximation and Projection (UMAP) dimensional reduction (RunUMAP). Markers for each cluster were identified with FindAllMarkers using default parameters, and cluster identity was determined based on the presence of known markers and populations previously described^34^. For cell-cell ligand-receptor interaction analysis, Seurat objects for acini were analyzed using CellChat^98^ with standard parameters. For lineage trajectory analysis, Seurat objects for acini were processed with Slingshot^40^, and then pseudotime was calculate with Monocle 3^99^.

### Ex vivo lineage tracing

Salivary glands from *Bmi1*^CreERT2^;Rosa26-RFP mice (recombination induced 24 hours prior) were mechanically dissected into < 1-mm pieces, placed in complete media (DMEM/F12 (Gibco, 11320033) + 50 μg/mL L-ascorbic acid (Sigma-Aldrich A4544) + 50 μg/mL holo-transferrin (Sigma-Aldrich, T1283) + 1% pen/strep (Gibco, 15140122)) in the presence or absence of 200 nM forskolin (Enzo Life Sciences, BML-CN100-0010), or DMSO (Thermo Fisher Scientific, BP231-100), and cultured for 48 hours before being fixed in 4% PFA for immunofluorescence analyses.

### In vivo lineage tracing

Lineage traced acinar cells was performed in healthy and IR *Bmi1*^CreERT^;Rosa26RFP mice as follows. Non-IR mice and IR mice (at 30 days post-IR) were injected i.p. with 2.5 mg tamoxifen per 1g body weight or vehicle and chased for 30 days. IR mice that were treated with saline or pilocarpine and non-IR mice were injected i.p. with 2.5 mg tamoxifen or vehicle per 10 g body weight at 91 days post-IR and chased for 30 days, with mice sacrificed at day 123 post-IR.

### Live cell imaging

Salivary gland explants or organoids were labeled with Fluo-4 NW Calcium Assay Kit (Invitrogen F36206) according to manufacturer’s protocol, BioTracker ATP-Red (Millipore SCT045) at 1:1000, Mitotracker Deep Red (Invitrogen M22426) at 300 nM, and/or with CellMask Green (Invitrogen C37608) at 1.5 µL/mL for 15 min at 37℃. After 3x brief rinses in HBBS, specimens mounted in glass-bottomed dishes (MatTek P35G-1.5-14-C) were imaged in 1xFluoroBrite DMEM (Gibco A1896701). For explants, wet filter paper and coverslip were used to secure the explants and prevent drifting. Imaging was performed at 37°C in an Okolab humidified microenvironmental chamber on the Nikon spinning disk confocal microscope. Identical laser power, exposure and gain settings were applied for all conditions within a set of experiments. For quantification of global trends, explants or organoids (n = 3-4) were labelled as above and imaged using CLARIOstar Plus microplate reader (BMG Labtech) at the respective wavelengths of each agent.

### Western Blotting and Analysis

Submandibular glands were lysed in RIPA buffer (Abcam, ab156034) supplemented with protease inhibitor cocktail (Cell Signaling Technology, 5872). Protein concentrations were measured by DC Protein Assay (Bio-Rad 5000116). Samples (∼60 µg lysates/lane) were resolved in NuPAGE 4-12% Bis-Tris gel (Invitrogen, NP0322BOX) at 150V for ∼1 hour at RT. Gels were transferred onto nitrocellulose membranes (Bio-Rad, 162-0177) using the iBlot transfer device (Invitrogen IB1001) for 7 min under 20 V. Membranes were blocked in Intercept TBS blocking buffer (LI-COR 927-60001) and incubated with primary antibodies (Supplementary Table S1) diluted in blocking buffer supplemented with 0.1% Tween-20 at 4℃ overnight. Signals were detected using IRDye secondary antibodies (LI-COR) and scanned on Odyssey CLx scanner (LI-COR, Model 9140). Band intensities were quantified using Image Studio Lite (LI-COR).

### Statistical Analysis

Statistical tests were performed using GraphPad Prism software v10. Data are plotted as individual data points with means ± SD. Groups with only two comparisons were analyzed with a two-tailed unpaired Student’s t-test. For multiple comparisons, an ordinary one-way analysis of variance (ANOVA) was used, followed by either a Tukey’s post-hoc comparisons test (for comparing means of multiple groups) or a Dunnett’s test. Significance was assessed using P value cutoffs.

### Data and materials availability

Bulk and snRNA-seq data have been deposited in the Gene Expression Omnibus database (https://ncbi.nlm.nih.gov/geo) under the accession no. GSE272766.

---

**Table S1:**
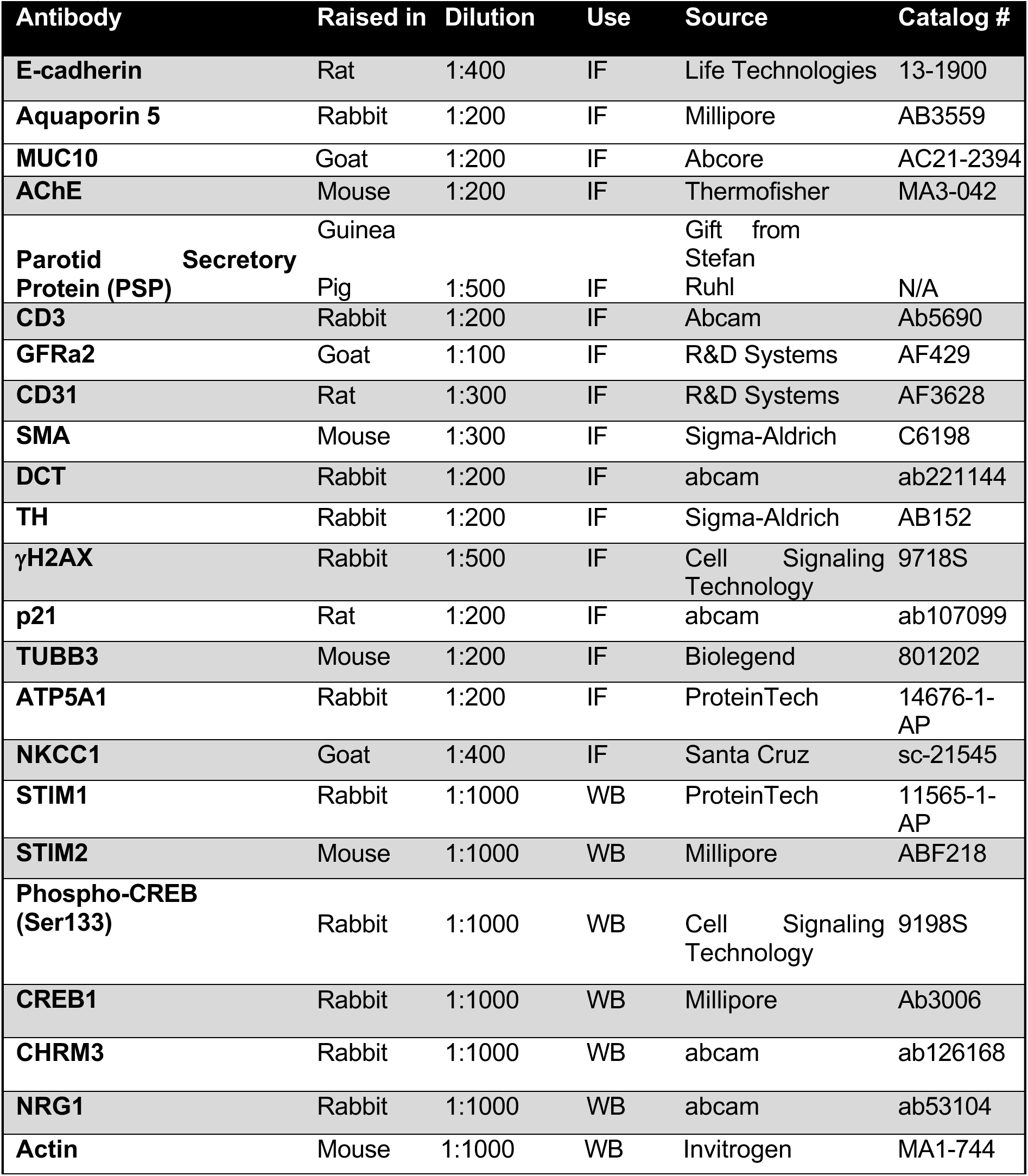
List of antibodies used.

## ACKNOWLEDGEMENTS

We thank Drs. Matthew Hoffman, Abigail Tucker, Susan Fisher and Jeffrey Bush for their comments on the manuscript. We thank Dr. Bjoern Schwer (University of California, San Francisco) for providing the γH2AX antibody and staining protocol. We thank UCSF Parnassus Flow Cytometry CoLab for the use of equipment and expertise, under the grants RRID: SSR 018206, DRC Center Grant NIH P30 DK063720, NIH S10 1S10OD021822-01 and NIH S10 1S10OD026940-01. The research was supported by NIDCR 1R35DE028255 and NIDCR CDOCTOR U24DE026914.

## AUTHOR CONTRIBUTIONS

Conceptualization: J.L., L.X.T., B.S., I.M.A.L., and S.M.K. Methodology: J.L., L.X.T., B.S., N.G., C.Y., M.H., C.T., Y.C.H, I.M.A.L, and S.M.K. Investigation: J.L., L.X.T., B.S., N.G., C.Y., S.V.N, L.B., N.C.P., S.M., H.S., Y.E., Y.T.C., L.A., E.G., M.H., C.T., Y.C.H, C.S.B., I.M.A.L, and S.M.K. Formal analysis: J.L., L.X.T., B.S., N.G., C.Y. Visualization: J.L., L.X.T., B.S., N.G., C.Y., I.M.A.L, and S.M.K. Data curation: J.L., I.M.A.L, and S.M.K. Project administration: S.M.K. Funding acquisition: S.M.K. Supervision: S.M.K. Writing—original draft: J.L., L.X.T., I.M.A.L, and S.M.K. Writing—review and editing: J.L., L.X.T., B.S., N.G., C.Y., C.S.B., I.M.A.L, and S.M.K.

## DECLARATION OF AI-ASSISTED TECHNOLOGIES

No AI-assisted technologies were used.

